# Heterotrophic succession of invertebrate assemblages on fallen guava fruits in a tropical grove

**DOI:** 10.64898/2026.10.05.756615

**Authors:** Sourabh Biswas, Silanjan Bhattacharyya

## Abstract

Fallen fleshy fruits are short-lived resource patches that can support diverse but transient invertebrate assemblages, yet their successional dynamics remain poorly documented in tropical human-modified landscapes. We examined heterotrophic succession on fallen ripe guava fruits, Psidium guajava, in a tropical grove in Barasat, West Bengal, India. Eight fruits were placed in exposed and shaded microhabitats and followed from fresh to degraded stages over 22 days. Invertebrates were recorded using repeated scan sampling, and fruit-day data were analysed for taxon richness, Shannon diversity, total peak relative abundance, community composition and stage-associated indicator taxa. Across 141 fruit-days, we recorded 44 invertebrate taxa. Taxon richness, Shannon diversity and total peak relative abundance were highest during fresh and pre-rotten stages and declined sharply during post-rotten and degraded stages. Decomposition stage was the strongest predictor of alpha diversity, resource-use intensity and community composition, whereas exposed–shaded microhabitat differences were weaker. Indicator taxa showed clear successional replacement: fresh fruits were associated with drosophilid and tephritid flies; fresh and pre-rotten fruits with nitidulid beetles and ants; and later stages with collembolans, staphylinids, blattodeans and other detrital-stage taxa. Ants, especially Pheidole sp., were conspicuous resource users and showed guarding, hoarding, predation and pulp or seed removal behaviour, although their effects on other taxa remain observational. These results show that fallen guava fruits function as ephemeral heterotrophic resource patches, supporting rapid invertebrate turnover at fine temporal scales in a tropical grove.

## Introduction

Short-lived organic resources occur as discrete and temporary patches within ecosystems and can support concentrated assemblages of consumers, decomposers, predators and scavengers. Carrion, dung, fungal fruiting bodies, decaying wood and decomposing plant material are examples of such ephemeral resource patches, whose availability and condition change rapidly through time (Barton et al. 2013; Butterworth, Benbow, and Barton 2022; Case et al. 2025). Because these resources are finite and progressively transformed or depleted, the communities exploiting them may change before a persistent equilibrium assemblage can develop. Ephemeral resource patches therefore provide useful systems for examining how colonization, resource use and community composition change over comparatively short temporal scales.

Fallen fleshy fruits represent a particularly dynamic form of ephemeral resource. After reaching the ground, fruits undergo progressive physical and biological changes that alter the conditions available to invertebrate consumers(Martins et al. 2026; Souza-Silva and Ferreira 2004). Gradual changes in the composition of invertebrate communities associated with such changing resources have been described as heterotrophic succession (Souza-silva and Ferreira 2004). Studies of *Alibertia edulis*, baobab (*Adansonia rubrostipa*) and *Psidium guajava* have documented differences in invertebrate richness, abundance or assemblage composition among fruit conditions or successional stages ((Łukasik and Johnson 2007; Silva and Ferreira 2011; Souza-silva and Ferreira 2004). In *P. guajava* specifically, physical and biological differences among fruit states have been interpreted as a heterotrophic successional gradient in which changing resource quality creates different opportunities for frugivorous invertebrates (Silva & Ferreira, 2011). Fallen fruits can also remain important resources for fruit-associated insects under conditions when other resources are limited (Deconninck et al. 2024). Together, these studies show that changes in fruit condition can be accompanied by substantial restructuring of associated invertebrate assemblages.

Much of this evidence, however, comes from comparisons among fruits representing different conditions or resource categories. Repeatedly following the same individual fallen-fruit patches from fresh condition through advanced decomposition provides a complementary way to resolve succession at finer temporal scales (Butterworth et al. 2022; Lettenmaier et al. 2025; Silva and Ferreira 2011). Such repeated observations can distinguish a simple decline in resource use from progressive replacement of taxa as a fruit passes through successive resource states. This distinction is especially relevant for short-lived resources, where substantial community change may occur within days or weeks rather than across the longer timescales commonly associated with ecological succession(Anderson 2007; Devánová et al. 2023; Purahong et al. 2016).

Although decomposition itself should impose a strong temporal gradient, the trajectory experienced by a fallen fruit may also depend on the microsite in which it decomposes. Exposed and shaded locations can differ in environmental conditions that potentially influence resource deterioration and invertebrate activity (Deconninck et al. 2024; Müller et al. 2020). Experimental works on another ephemeral decomposing resource has shown that environmental conditions can alter successional trajectories and beetle community composition during dead-wood decay (Sbaraglia et al. 2025; Vindstad et al. 2020). Exposed–shaded differences in fallen-fruit assemblages could therefore arise through differences in the rate or pathway of fruit degradation (Martins et al. 2026; De Paz, Gobbi, and Raffaele 2018). At the same time, such microsites may also differ in colonization opportunities and local biotic interactions (Sbaraglia et al. 2025; Vindstad et al. 2020). Without direct measurements of the underlying mechanisms, exposed–shaded contrasts are consequently best regarded as microhabitat-associated variation that may modify the broader decomposition trajectory (Kim et al. 2021; Vindstad et al. 2020).

Biotic interactions within the resource patch may provide an additional source of variation. Ants are particularly relevant because they can rapidly discover and remove concentrated resources; in tropical forests they can account for a major proportion of experimentally available resource removal (Griffiths et al. 2017). Ants also interact with fruits and seeds lacking specialized adaptations for ant dispersal, including through pulp exploitation and secondary diaspore removal (Anjos et al. 2020; Bieber et al. 2014; Clemente and Whitehead 2019; Pizo and Oliveira 2000). On fallen fruits, abundant ants may therefore act not only as consumers but also as potential modifiers of resource accessibility through removal, guarding or interactions with other invertebrates (Bieber et al. 2014; King et al. 2018; Morris et al. 2026). However, in the absence of experimental ant exclusion, relationships between ant abundance and the wider fruit-associated assemblage can demonstrate association but not causal suppression (Griffiths et al. 2017; Kim et al. 2021; Lunde et al. 2022).

Here, we followed invertebrate assemblages exploiting standardized fallen ripe guava fruits, *Psidium guajava*, in a tropical grove in Barasat, West Bengal, India, from fresh condition through advanced degradation. Fruits were placed in exposed and shaded microhabitats and repeatedly observed throughout decomposition. We asked (1) whether taxon richness, Shannon diversity and total peak relative abundance changed across decomposition stages; (2) whether community composition and the identities of stage-associated taxa changed as fruits decomposed; (3) whether these patterns differed between exposed and shaded microhabitats; and (4) whether the abundance of the dominant ant *Pheidole*sp. 1 was associated with variation in the non-ant assemblage. We predicted that decomposition stage would represent the principal axis of assemblage change, with richness and peak relative abundance remaining highest during fresh and intermediate stages before declining as fruits became post-rotten and degraded. We expected microhabitat-associated differences to be smaller than the temporal effect of decomposition. Given our observations of conspicuous *Pheidole*activity, we additionally explored whether increasing *Pheidole* abundance was associated with reduced non-ant richness or peak relative abundance, while treating these relationships as observational rather than causal.

## Materials and methods

### Study site

The study was conducted at Barasat Kisoloy, Barasat, West Bengal, India (22.7174°N, 88.4805°E), a tropical grove embedded within a human-modified landscape (Figure S1). The grove contains secondary-growth vegetation, scattered trees, leaf litter, open patches, shaded vegetation and two water bodies. Field observations were conducted from February to June 2016, covering late winter, summer and the pre-monsoon period. Observations continued until invertebrate activity on each fruit had ceased or the fruit had reached a degraded condition.

### Experimental fruit patches and microhabitat classification

Ripe guava fruits, *Psidium guajava*, were used as standardized ephemeral resource patches. Fruits were collected from nearby local markets, and fruits of approximately similar size and weight were used as far as possible. Fruit weight was recorded before placement and later included as a covariate in the statistical analyses. Each fruit represented one experimental resource patch and was followed through time from placement to degradation.

To approximate the impact damage experienced by naturally fallen ripe fruits, each guava was manually tossed onto the ground within its assigned microhabitat. This punctured or broke the fruit surface and exposed the pulp, creating access points similar to those observed in naturally fallen fruits. Fruits were placed on the ground over leaf litter and were not subsequently manipulated except during observation or specimen collection for identification.

Two microhabitat types were selected: exposed and shaded. Exposed microhabitats were relatively open areas where fruits received direct or substantial sunlight during the day. Shaded microhabitats were under dense canopy or bush cover, where direct sunlight was limited or largely absent. A total of eight fruits were used, with four fruits placed in exposed microhabitats and four in shaded microhabitats. Fruit identity was used as the replicate unit in subsequent analyses.

### Decomposition stages

Fruit decomposition was classified into four stages based on time since placement, external colour, pulp condition and odour. The fresh stage corresponded approximately to days 1–3, when the exocarp remained yellowish, the pulp was pale and no strong rotten odour was present. The pre-rotten stage corresponded approximately to days 4–10, when the exocarp began turning brown, the pulp became slightly brownish and rotten odour was detectable. The post-rotten stage corresponded approximately to days 11–18, when the exocarp became dark brown, the pulp was reduced and strongly decomposed, and odour was pronounced. The degraded stage corresponded approximately to days 19–22, when most pulp had disappeared and only hard exocarp or dried remains persisted. These stages were used to represent the temporal progression of heterotrophic succession on the fallen fruit resource.

### Invertebrate observations

Invertebrates visiting or exploiting the guava fruits were recorded using instantaneous scan sampling. Daylight observation periods were divided into four broad sessions: morning, noon, afternoon and evening. During each observation day, each fruit was scanned at hourly intervals, and all visible invertebrate taxa associated with the fruit were recorded. Ambient temperature and relative humidity were recorded during field observations and included as covariates in the statistical analyses.

For each fruit, day and taxon, the highest number of individuals observed in any hourly scan was retained as the daily maximum scan abundance. This value was treated as an index of peak relative abundance or maximum daily use of the fruit resource, not as true total daily abundance. Presence or absence of each taxon on each fruit-day was also derived from these scan observations.

Qualitative interaction data were recorded during scan sampling and direct observation. These included coexistence among taxa, aggressive encounters, apparent predation, ant guarding or hoarding behaviour, and seed or pulp removal. Observations were made from a distance that minimized disturbance, and selected behaviours were photographed or video-recorded when possible.

## Specimen collection and identification

Invertebrates were identified using direct field observations, photographic records and collected specimens. Visually distinctive taxa were identified in the field or from photographs using available taxonomic resources and expert consultation. Invertebrates requiring closer examination, including small Coleoptera, Hymenoptera, Diptera, Hemiptera and other arthropods, were collected using hand picking, forceps, brushes or sweep nets depending on size and mobility. Small specimens were preserved in 70% ethanol.

Specimen collection was conducted primarily for taxonomic identification and was not used for abundance estimation. Specimens were submitted to the Zoological Survey of India, Kolkata, for identification. Taxa that could not be identified to species level were retained as morphospecies or higher taxonomic units and used consistently across the dataset.

### Seed and pulp removal by ants

Ant-mediated removal of guava pulp and seeds was recorded opportunistically during behavioural observations. When ants were observed carrying seeds or pulp fragments away from the fruit, the direction and distance of removal were measured. The straight-line distance between the fruit and the destination point or nest entrance was estimated using thread and later measured using measuring tape or a centimetre scale. These observations were used as natural-history evidence of ant participation in pulp or seed removal, not as a formal test of seed dispersal.

### Statistical analysis

All analyses were conducted in R (R Core Team 2022). Data preparation used the tidyverse (Wickham et al. 2019). The fruit-day was the sampling unit for taxon richness, Shannon diversity and total peak relative abundance. Because fruits were observed repeatedly, fruit identity was included as a random intercept in mixed models. Microhabitat and decomposition stage were categorical predictors, with exposed microhabitat and fresh fruit as reference categories. Temperature, humidity and fruit weight were standardized and included as covariates in the main mixed models. Taxon richness was the number of taxa with positive peak relative abundance on each fruit-day. Shannon diversity was calculated from the taxon-level peak relative abundance matrix using vegan (Oksanen, 2015). Richness was analysed with Poisson and negative-binomial generalized linear mixed models fitted using glmmTMB (Bolker 2019). Model residuals and dispersion were examined using DHARMa (Hartig and Hartig 2022). Shannon diversity was analysed with a linear mixed model fitted using lme4 (Bates et al. 2015), with fixed-effect tests obtained using lmerTest (Kuznetsova, Brockhoff, and Christensen 2017).

Total peak relative abundance was the sum of the maximum daily scan count for each taxon within a fruit-day. It was analysed separately from richness as an index of peak resource use, rather than a count of all individuals observed during the day. Poisson, negative-binomial and COM-Poisson mixed models were fitted using glmmTMB and assessed using information criteria, convergence checks and DHARMa residual diagnostics. Estimated marginal means and pairwise comparisons were obtained using emmeans (Lenth 2023); Tukey adjustment was applied to the reported comparisons for total peak relative abundance.

Community composition was analysed with vegan using two approaches. For abundance-weighted analyses, peak relative abundances were square-root transformed before calculating Bray–Curtis dissimilarities. For occurrence-based analyses, the matrix was converted to presence/absence data before calculating Jaccard dissimilarities. Fruit-days with no recorded taxa were excluded. Non-metric multidimensional scaling was used to visualize composition.

Permutational multivariate analysis of variance (vegan, adonis2; 999 permutations) examined decomposition stage, microhabitat and their interaction. The full interaction models used unrestricted permutations and were interpreted as exploratory because fruit-days from the same fruit were repeated observations. Stage-only tests and additive stage-plus-microhabitat models used permutations constrained within fruit identity; these provide the more appropriate permutation tests for changes in stage within fruits. Bray–Curtis multivariate dispersion was assessed separately by stage and microhabitat using betadisper and permutation tests in vegan.

Indicator taxon analysis used multipatt in indicspecies (Cáceres and Legendre 2009) with the IndVal.g statistic and 999 permutations specified through permute (Simpson et al. 2019). Presence/absence analysis was treated as the primary result, with analysis of square-root-transformed peak relative abundance as supporting evidence. Taxa with unadjusted permutation p values below 0.05 were identified as potential indicators. These tests were interpreted exploratorily because their permutations were not constrained by fruit identity.

Exploratory mixed models fitted using glmmTMB tested associations between the peak relative abundance of Pheidole taxa and non-ant richness or non-ant peak relative abundance. Models accounted for microhabitat, decomposition stage, temperature, humidity and repeated observations of fruit identity. These observational associations were not interpreted as evidence that ants suppressed other taxa.

## Results

### Dataset overview

After filtering the dataset to retain records marked for analysis, the final dataset contained 7,744 taxon-level observations from eight experimental fruits followed across days 1– 22 of decomposition. In total, 44 invertebrate taxa were recorded, including ants, flies, beetles, hemipterans, collembolans, blattodeans, millipedes, spiders, mites and butterflies (Table S1). The sample-level dataset contained 141 fruit-days, including 67 exposed and 74 shaded fruit-days. Across decomposition stages, there were 24 fresh, 48 pre-rotten, 54 post-rotten and 15 degraded fruit-days. Two fruit-days with missing humidity values were excluded from mixed models that included humidity as a covariate, giving a model sample size of 139 fruit-days. Community-composition analyses excluded fruit-days with no recorded taxa, leaving 124 fruit-days for Bray–Curtis and Jaccard analyses. Full dataset structure and analysis-specific sample sizes are provided in Table S2.

### Alpha diversity and resource-use intensity declined during decomposition

Taxon richness, Shannon diversity and total peak relative abundance varied strongly across fruit decomposition stages (Fig. 1; Table S2). Descriptively, richness was highest during the fresh and pre-rotten stages and declined during the post-rotten and degraded stages (Table 1). Exposed fruits declined from 11.33 ± 1.28 SE taxa per fruit-day in the fresh stage to 3.67 ± 1.96 SE in the degraded stage. Shaded fruits showed a similar decline, from 8.17 ± 0.92 SE taxa per fruit-day in the fresh stage to 0.89 ± 0.77 SE in the degraded stage. Shannon diversity followed the same broad pattern, declining from 1.49 ± 0.17 SE in fresh exposed fruits and 1.53 ± 0.14 SE in fresh shaded fruits to 0.20 ± 0.20 SE and 0.16 ± 0.16 SE, respectively, in degraded fruits.

**Table 1.**
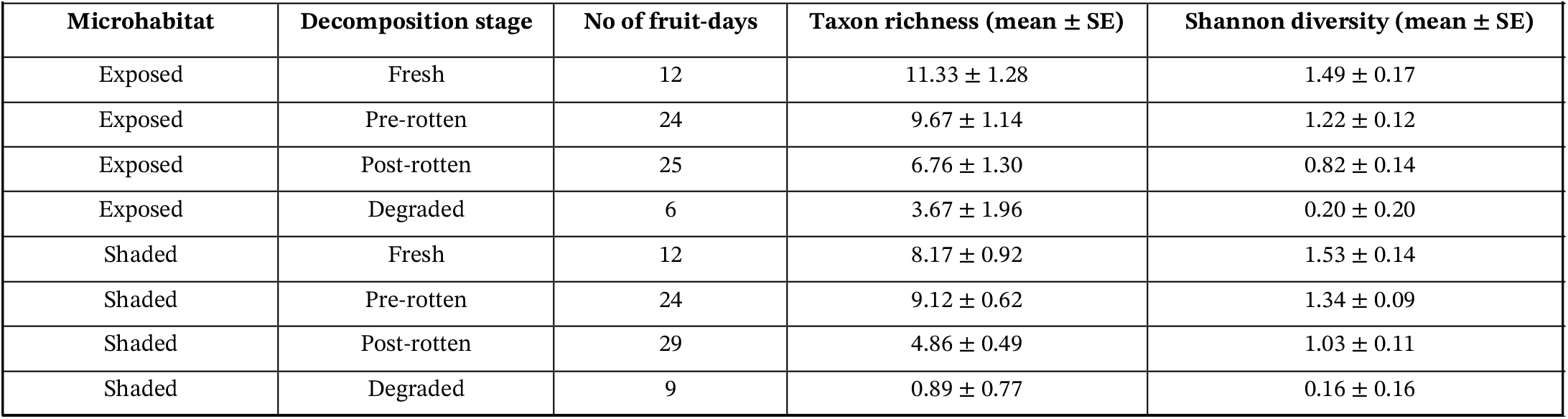
Sample-level alpha diversity across decomposition stages and microhabitats. Values are presented as mean ± SE. Taxon richness represents the number of taxa recorded per fruit-day, and Shannon diversity was calculated using peak relative abundance

| Microhabitat | Decomposition stage | No of fruit-days | Taxon richness (mean ± SE) | Shannon diversity (mean ± SE) |
| --- | --- | --- | --- | --- |
| Exposed | Fresh | 12 | $11.33 \pm 1.28$ | $1.49 \pm 0.17$ |
| Exposed | Pre-rotten | 24 | $9.67 \pm 1.14$ | $1.22 \pm 0.12$ |
| Exposed | Post-rotten | 25 | $6.76 \pm 1.30$ | $0.82 \pm 0.14$ |
| Exposed | Degraded | 6 | $3.67 \pm 1.96$ | $0.20 \pm 0.20$ |
| Shaded | Fresh | 12 | $8.17 \pm 0.92$ | $1.53 \pm 0.14$ |
| Shaded | Pre-rotten | 24 | $9.12 \pm 0.62$ | $1.34 \pm 0.09$ |
| Shaded | Post-rotten | 29 | $4.86 \pm 0.49$ | $1.03 \pm 0.11$ |
| Shaded | Degraded | 9 | $0.89 \pm 0.77$ | $0.16 \pm 0.16$ |

**Figure 1.**
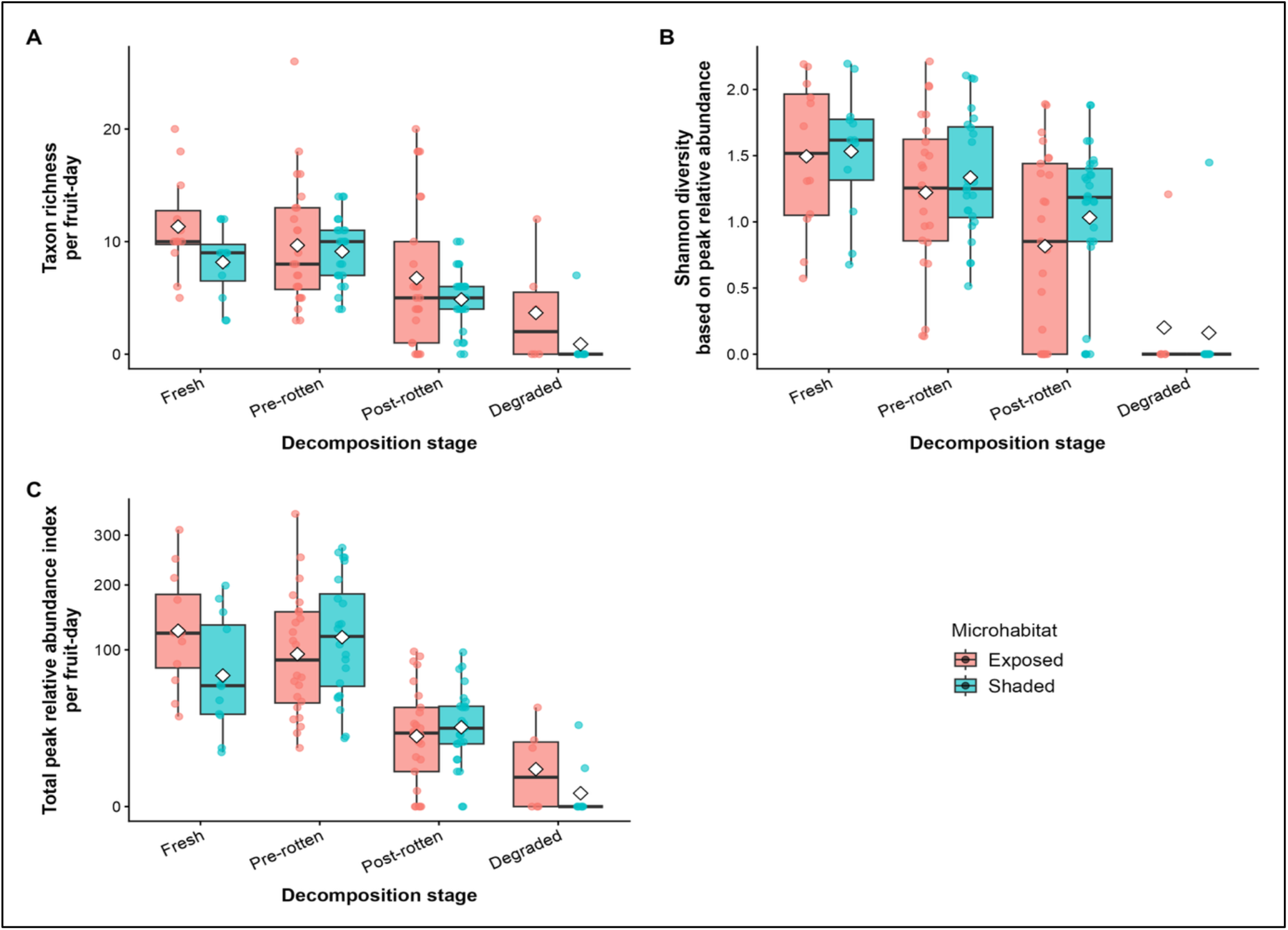
Alpha diversity and total peak relative abundance of invertebrates across guava fruit decomposition stages. Changes in (A) taxon richness per fruit-day, (B) Shannon diversity calculated from taxon-level peak relative abundance, and (C) total peak relative abundance index on fallen ripe Psidium guajava fruits across decomposition stages in exposed and shaded microhabitats. Boxplots show the distribution of fruit-day values, points represent individual fruit-days, and white diamonds indicate mean values. Richness, Shannon diversity and total peak relative abundance were highest during the fresh and pre-rotten stages and declined sharply during the post-rotten and degraded stages, indicating reduced diversity and resource-use intensity as fruit decomposition progressed.

Taxon richness was analysed using a negative-binomial GLMM. Decomposition stage was the strongest predictor of richness (Table S3). Relative to fresh fruits, richness was significantly lower in post-rotten fruits (z = −3.471, p < 0.001) and degraded fruits (z = −5.312, p < 0.001), whereas pre-rotten fruits did not differ from fresh fruits (z = −0.061, p = 0.951).

Shaded fruits did not differ significantly from exposed fruits (z = −0.965, p = 0.335), and the microhabitat × decomposition stage interaction was not supported. Model-estimated richness was highest in fresh and pre-rotten fruits and lowest in degraded fruits (Table S3B). Pairwise comparisons showed no significant difference between fresh and pre-rotten fruits, but both early stages had significantly higher richness than post-rotten and degraded fruits(Table S3C).

Shannon diversity showed a parallel decline. The Shannon diversity mixed model indicated that decomposition stage accounted for more variation than microhabitat or the microhabitat × stage interaction (Table S4A). Model-estimated Shannon diversity was highest in fresh fruits and lowest in degraded fruits, declining from approximately 1.46–1.49 in fresh fruits to 0.05–0.13 in degraded fruits across microhabitats (Table S4B-C).

Total peak relative abundance was analysed separately from richness because it represents maximum daily resource-use intensity rather than total abundance. The accepted COM-Poisson GLMM showed that decomposition stage was again the strongest predictor (Figure 1, Table S5). Relative to fresh fruits, total peak relative abundance did not differ significantly in the pre-rotten stage (z = −0.687, p = 0.492), but was significantly lower in post-rotten fruits (z = −6.522, p < 0.001) and degraded fruits (z = −6.323, p < 0.001). Shaded fruits did not differ significantly from exposed fruits overall (z = −1.358, p = 0.174), and temperature, humidity and fruit weight were not significant predictors.

One significant interaction was detected between shaded microhabitat and the post-rotten stage (z = 2.237, p < 0.05; Table S5), indicating that the shape of the decline from fresh to post-rotten fruits differed between microhabitats. However, direct exposed–shaded contrasts within individual decomposition stages were not significant. Therefore, this interaction is best interpreted as a difference in the temporal pattern of decline rather than strong evidence for a microhabitat effect at any single stage. Overall, richness, Shannon diversity and total peak relative abundance remained high during fresh and pre-rotten stages and declined sharply after the transition to post-rotten and degraded stages (Fig. 1).

### Broad taxonomic composition changed through decomposition

Order-level summaries showed that the decline in richness and resource-use intensity was accompanied by broad taxonomic turnover (Fig. 2; Fig. S2). Fresh and pre-rotten fruits were dominated mainly by Hymenoptera, Diptera and Coleoptera, although their relative contributions differed between exposed and shaded fruits. In exposed fresh fruits, Hymenoptera contributed the largest share of mean peak relative abundance, whereas in shaded fresh fruits Diptera contributed the largest share. During the pre-rotten stage, Hymenoptera became the dominant order in both microhabitats, with Coleoptera also contributing substantially.

**Figure 2.**
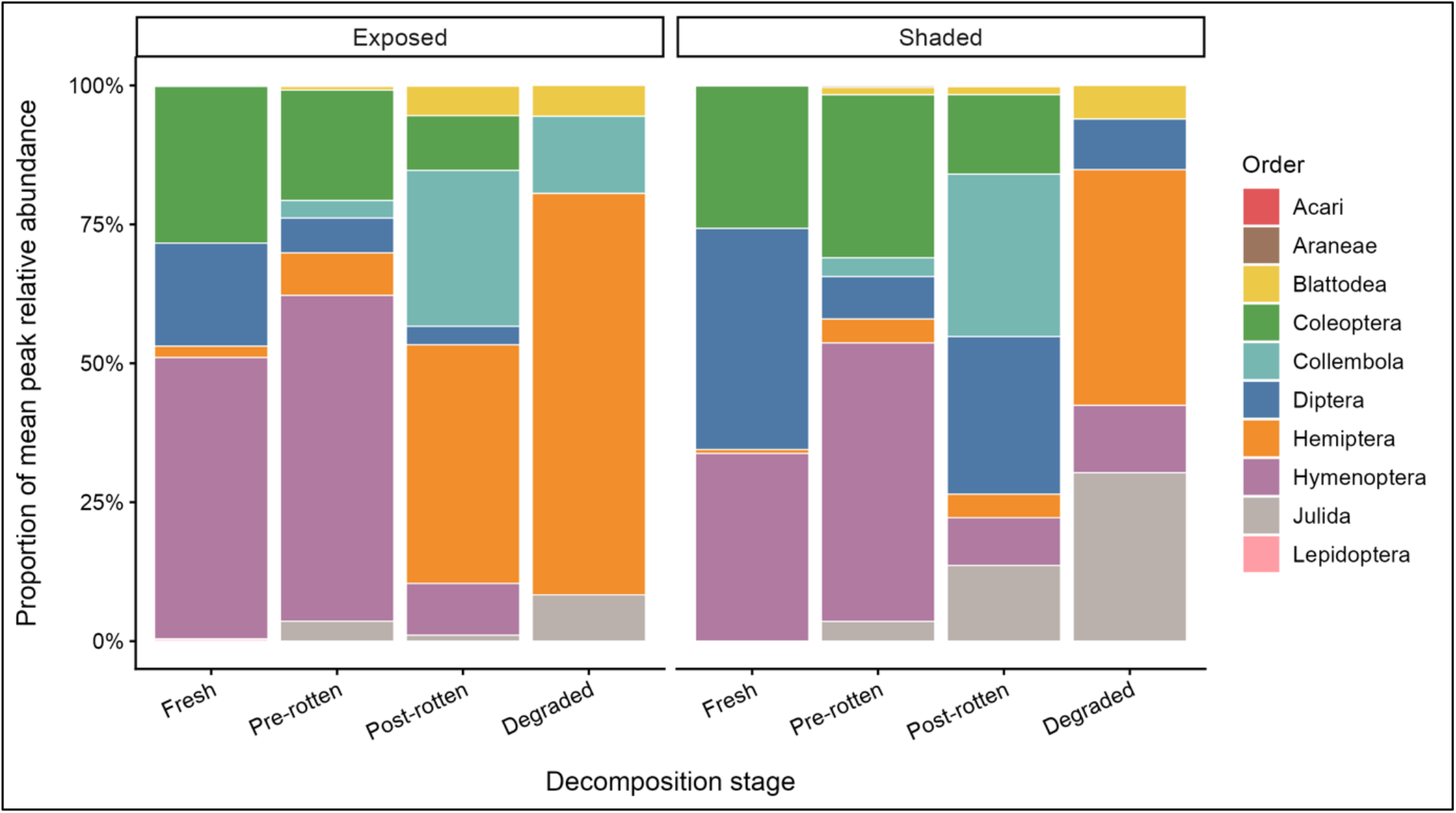
Order-level composition of invertebrate assemblages across guava fruit decomposition stages. Proportional contribution of major invertebrate orders to mean peak relative abundance on fallen ripe Psidium guajava fruits across decomposition stages in exposed and shaded microhabitats. Each stacked bar represents the relative contribution of orders within a given decomposition stage and microhabitat. Fresh and pre-rotten fruits were dominated mainly by Hymenoptera, Diptera and Coleoptera, whereas post-rotten and degraded fruits showed increased proportional contributions from Hemiptera, Collembola, Julida and other later-stage groups. Because the degraded stage had low total peak relative abundance, proportional dominance in this stage should be interpreted cautiously.

Later stages showed a shift away from early dominance by Hymenoptera, Diptera and Coleoptera. Post-rotten and degraded fruits showed increased proportional contributions from Hemiptera, Collembola, Julida and Blattodea (Fig. 2; Fig. S2). Because degraded fruit-days had low total activity, high proportional contributions by individual orders in this stage should be interpreted cautiously (Fig. S3). Overall, the order-level results indicate that fruit decomposition involved both a decline in overall activity and a shift in broad taxonomic structure.

### Community composition changed across decomposition stages

Community composition differed among decomposition stages in both abundance-weighted and presence/absence analyses. NMDS ordination based on Bray–Curtis dissimilarity showed stage-associated separation among fruit-days, although ordination stress was moderate (stress = 0.227; Fig. 3). Therefore, the ordination should be interpreted as a broad visual summary rather than an exact representation of all pairwise distances. The ordination was nevertheless consistent with the alpha-diversity and order-level results: fresh and pre-rotten fruit-days partly overlapped, whereas post-rotten and degraded fruit-days shifted toward more compositionally distinct and reduced regions of ordination space. The presence/absence-based Jaccard NMDS showed a similar broad pattern and is provided in Fig. S4.

**Figure 3.**
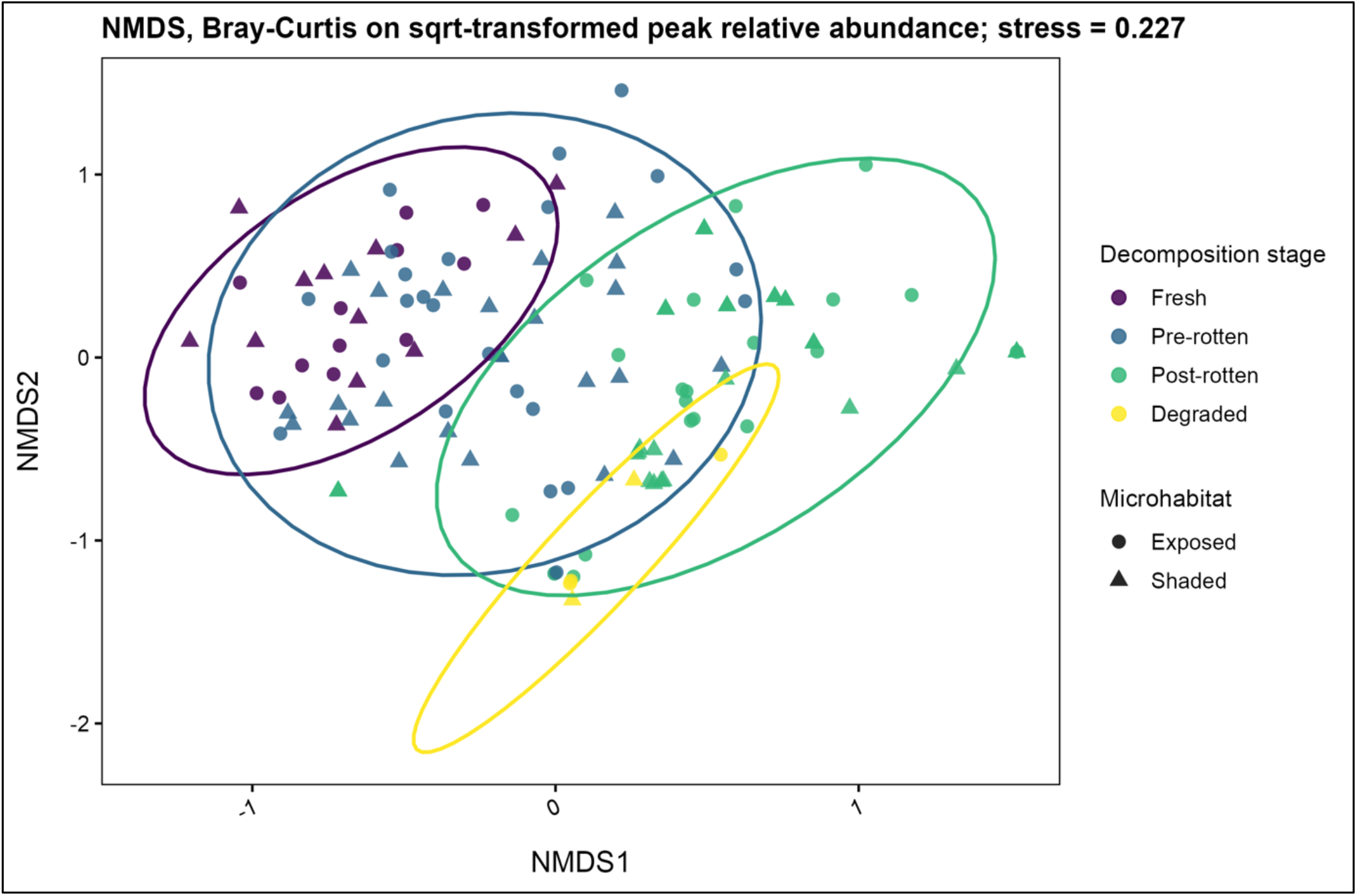
Abundance-weighted community composition of invertebrates across guava fruit decomposition stages. Non-metric multidimensional scaling (NMDS) ordination based on Bray–Curtis dissimilarity calculated from square-root-transformed peak relative abundance of invertebrate taxa recorded on fallen ripe *Psidium guajava* fruit-days. Points represent individual fruit-days, colours indicate decomposition stage, and shapes indicate microhabitat type: exposed or shaded. Ellipses summarize the dispersion of fruit-days within each decomposition stage. The ordination shows stage-associated turnover in abundance-weighted community composition, with fresh and pre-rotten fruit-days partly overlapping and post-rotten and degraded fruit-days shifting toward a more distinct region of ordination space. The moderate stress value (stress = 0.227) indicates that the plot should be interpreted as a broad visual summary of compositional differences rather than an exact representation of all pairwise dissimilarities.

Restricted PERMANOVA, with permutations constrained within fruit identity, showed that decomposition stage significantly explained abundance-weighted community composition based on Bray–Curtis dissimilarity (R^2^ = 0.1849, F = 9.074, p = 0.001; Table 1; Table S6A-D). Presence/absence analysis gave a similar result, with decomposition stage significantly explaining Jaccard community composition (R^2^ = 0.1407, F = 6.550, p = 0.001; Table 1; Table S6A-D). Thus, decomposition stage accounted for approximately 18.5% of variation in abundance-weighted composition and 14.1% of variation in taxon occurrence.

The full exploratory PERMANOVA including the microhabitat × decomposition stage interaction did not support an interaction effect for either Bray–Curtis or Jaccard composition (Table S6A). An additive PERMANOVA showed that microhabitat had a statistically detectable but much smaller effect than decomposition stage (Table 2; Table S8). microhabitat explained 2.5% of variation in Bray–Curtis composition and 1.9% of variation in Jaccard composition, compared with 18.5% and 14.1%, respectively, for decomposition stage.

**Table 2.** Summary of PERMANOVA results testing effects of decomposition stage and microhabitat on fruit-associated invertebrate community composition. Bray–Curtis dissimilarity was calculated using square-root-transformed peak relative abundance, whereas Jaccard dissimilarity was calculated using presence/absence data. Restricted PERMANOVA tests used permutations constrained within fruit identity to account for repeated sampling of the same fruits through decomposition.

| Analysis | Term | Dissimilarity | R<sup>2</sup> | F | p | Permutation |
| --- | --- | --- | --- | --- | --- | --- |
| Restricted PERMANOVA | Decomposition stage | Bray–Curtis | 0.1849 | 9.074 | 0.001 | within fruit |
| Restricted PERMANOVA | Decomposition stage | Jaccard | 0.1407 | 6.550 | 0.001 | within fruit |
| Additive PERMANOVA | Microhabitat | Bray–Curtis | 0.025 | 3.739 | 0.001 | within fruit |
| Additive PERMANOVA | Microhabitat | Jaccard | 0.019 | 2.708 | 0.001 | within fruit |

Therefore, exposed and shaded fruits differed slightly in community composition, but decomposition stage was the dominant driver of community turnover.

Multivariate dispersion differed significantly among decomposition stages for Bray–Curtis dissimilarity (F = 12.539, p = 0.001), but not between microhabitats (F = 0.056, p = 0.823; Table S6C-D). Pairwise dispersion comparisons showed that degraded fruit-days had significantly lower dispersion than fresh, pre-rotten and post-rotten fruit-days (Table S7). The stage-level dispersion effect indicates that decomposition-stage differences reflected both shifts in community centroids and changes in within-stage variability (Table S8). Degraded fruit-days were compositionally more constrained, consistent with their low richness and low peak relative abundance.

### Indicator taxa identified stage-associated assemblage shifts

Indicator taxon analysis identified clear stage-associated changes in the invertebrate assemblage. Presence/absence analysis, treated as the primary conservative result, identified 14 significant indicator taxa out of 44 taxa (Table 3; Table S9). Fresh fruits were characterized by drosophilid and tephritid flies, including *Drosophila* sp. 1, *Drosophila ananassae*and *Bactrocera dorsalis*, as well as the ant *Ectomomyrmex annamitus*. These taxa represent early-stage indicators based on their occurrence patterns.

**Table 3.** Significant indicator taxa associated with decomposition stages of fallen guava fruits based on presence/absence data. Indicator taxa were identified using multilevel pattern analysis with the IndVal.g association function and 999 permutations. Only taxa significant at p < 0.05 are shown. Higher IndVal values indicate stronger association with the corresponding decomposition stage or stage combination.

| Taxon | Associated decomposition stage | IndVal | p-value | Ecological interpretation |
| --- | --- | --- | --- | --- |
| *Drosophila* sp.1 | Fresh | 0.740 | 0.001 | Early-stage indicator |
| *Drosophila ananassae* | Fresh | 0.535 | 0.031 | Early-stage indicator |
| *Bactrocera dorsalis* | Fresh | 0.482 | 0.019 | Early-stage indicator |
| *Ectomomyrmex annamitus* | Fresh | 0.386 | 0.049 | Early-stage indicator |
| *Epuraea* sp.1 | Fresh + Pre-rotten | 0.700 | 0.002 | Early-to-intermediate stage indicator |
| *Pheidole* sp.1 | Fresh + Pre-rotten | 0.683 | 0.002 | Early-to-intermediate stage indicator |
| *Urophorus* sp. | Fresh + Pre-rotten | 0.669 | 0.001 | Early-to-intermediate stage indicator |
| *Epuraea* sp.2 | Fresh + Pre-rotten | 0.646 | 0.004 | Early-to-intermediate stage indicator |
| *Camponotus crassisquamis* | Fresh + Pre-rotten | 0.606 | 0.025 | Early-to-intermediate stage indicator |
| *Lasicodactylus* sp. | Fresh + Pre-rotten | 0.601 | 0.003 | Early-to-intermediate stage indicator |
| *Drosophila* sp. | Fresh + Pre-rotten | 0.471 | 0.035 | Early-to-intermediate stage indicator |
| Unidentified Staphylinidae sp. | Pre-rotten + Post-rotten | 0.682 | 0.005 | Mid-successional indicator |
| Unidentified Collembola sp.2 | Post-rotten + Degraded | 0.665 | 0.003 | Late-stage indicator |
| Unidentified Blattodea sp.2 | Pre-rotten + Post-rotten + Degraded | 0.548 | 0.027 | Post-fresh decomposition indicator |

Several taxa were associated with the combined fresh and pre-rotten stages, indicating that early-to-intermediate decomposition supported a distinct assemblage (Table 2; Table S9). These included nitidulid beetles such as *Epuraea* sp. 1, *Epuraea* sp. 2, *Urophorus* sp. and *Lasicodactylus* sp., as well as ants including *Pheidole* sp. 1 and *Camponotus crassisquamis*. This period therefore represented the main phase of active resource use by fruit-associated flies, nitidulid beetles and ants.

Later decomposition stages were associated with different taxa (Table 2; Table S9). Unidentified Staphylinidae sp. was associated with the pre-rotten and post-rotten stages, unidentified Collembola sp. 2 with the post-rotten and degraded stages, and unidentified Blattodea sp. 2 with the broader pre-rotten, post-rotten and degraded stage combination. These patterns indicate a shift from early fruit-feeding and fruit-attracted taxa toward taxa more frequently associated with later decompositional or detrital conditions.

The abundance-weighted indicator analysis produced a similar pattern and is reported in Table S10. In that analysis, *Metochus uniguttatus* was additionally identified as a degraded-stage indicator. The close agreement between presence/absence and abundance-weighted analyses strengthens the interpretation that decomposition stage was associated with consistent taxon replacement. Dominant-taxon summaries also supported this pattern: the most abundant taxa overall included *Pheidole* sp. 1, *Solenopsis geminata, Metochus uniguttatus*, unidentified Staphylinidae sp., *Epuraea*sp. 1, unidentified Collembola sp. 2 and *Drosophila* sp. 1. These taxa contributed strongly to the stage- and microhabitat-specific abundance patterns shown in the heatmap of the top 20 taxa (Fig. 4).

**Figure 4.**
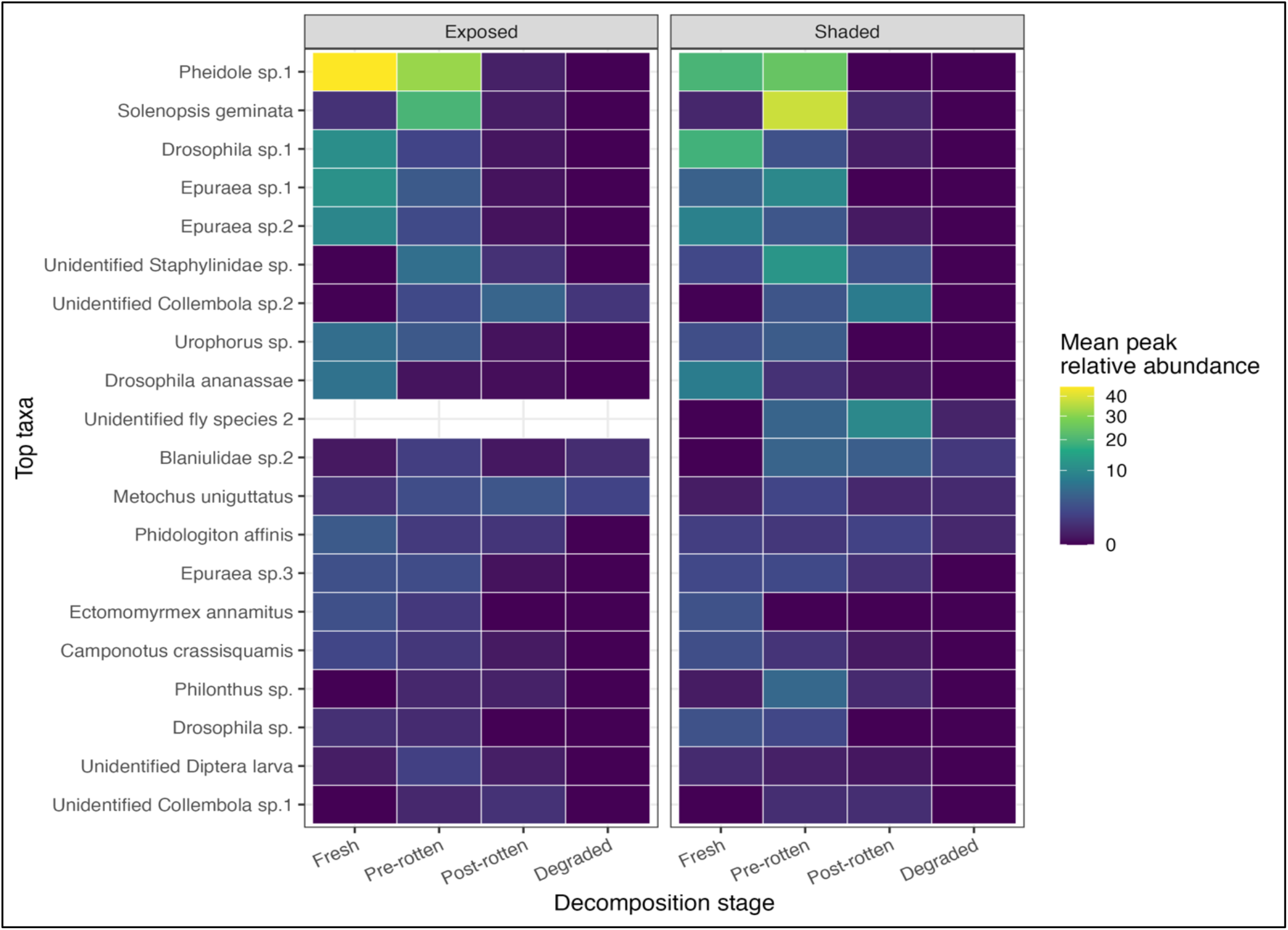
Stage- and microhabitat-specific abundance patterns of the most abundant taxa on fallen guava fruits. Heatmap showing mean peak relative abundance of the 20 most abundant invertebrate taxa recorded on fallen ripe Psidium guajava fruits across decomposition stages in exposed and shaded microhabitats. Colour intensity represents mean peak relative abundance within each stage–microhabitat combination, with brighter colours indicating higher values. The heatmap shows that several dominant taxa, including Pheidole sp.1, Solenopsis geminata, Drosophila sp.1 and Epuraea spp., were most abundant during fresh and pre-rotten stages, whereas taxa such as unidentified Collembola, unidentified Staphylinidae and Metochus uniguttatus were more associated with later decomposition stages. The figure illustrates taxon-level replacement during guava fruit decomposition.

### Natural-history observations and ant-associated patterns

Direct observations showed that fallen guava fruits functioned not only as feeding substrates but also as sites of behavioural interaction. Ants were among the most conspicuous users of the fruit resource. The commonest ant behaviour observed was feeding on fruit juice from punctured or exposed pulp. *Camponotus crassisquamis* responded early during the fresh stage and was observed carrying pulp fragments containing seeds away from the fruit. *Pheidole* sp. 1 also arrived during the fresh stage, but seed and pulp removal by this taxon was observed mainly during the pre-rotten and post-rotten stages. Observed removal distances ranged from 0.035 m to 8.37 m, with *Pheidole* sp. 1 recorded as the longest-distance remover.

*Pheidole* sp. 1 also showed conspicuous guarding or hoarding behaviour. Workers deposited soil particles or dry leaf fragments over the punctured surface of the fruit and later within the pulp. This behaviour partly covered the exposed pulp and appeared to restrict access to the fruit surface. Workers also guarded the fruit and showed aggression toward other invertebrates attempting to enter or exploit the resource. *Pheidole* sp. 1 was observed attacking or carrying maggots and some coleopterans, whereas Diptera and Coleoptera often co-occurred during the fresh stage and appeared to use different parts of the fruit.

Exploratory models provided only weak support for negative associations between *Pheidole* sp. 1 and the non-ant assemblage after accounting for decomposition stage and other covariates (Table S11; Table S12). *Pheidole* sp. 1 peak relative abundance showed a weak negative but non-significant association with non-ant richness (z = −1.681, p = 0.0929; Table S11; Fig. S5) and a marginal negative association with non-ant peak relative abundance (z = −1.905, p = 0.0568; Table S12; Fig. S6). In both models, decomposition stage remained the strongest predictor of non-ant richness and abundance. Group-specific models for Diptera, Coleoptera and maggots or larvae did not show significant negative associations with *Pheidole* sp. 1 abundance and are reported in Tables S13–S15. These observations suggest that dominant ants may modify access to fallen fruit patches, but because the analyses were observational and no ant-exclusion experiment was conducted, these patterns should be interpreted as natural-history evidence and associations rather than causal suppression.

## Discussion

Fallen ripe guava fruits supported a rapid and directional turnover of invertebrate assemblages over the 22-day decomposition period. Decomposition stage was consistently the strongest predictor of taxon richness, Shannon diversity, total peak relative abundance and community composition.

Richness and resource-use intensity remained high during the fresh and pre-rotten stages but declined markedly once fruits entered the post-rotten and degraded stages. At the same time, indicator taxa changed across decomposition, showing that the assemblage did not simply lose taxa as the resource was depleted. Instead, early fruit-associated taxa were progressively replaced by taxa characteristic of later decomposition stages. Taken together, these patterns are consistent with heterotrophic succession on an ephemeral resource patch, in which temporal change in the condition of a finite resource is accompanied by restructuring of the consumer assemblage.

This interpretation fits the broader ephemeral-resource framework, in which discrete resources change rapidly in condition and availability and may disappear before their associated communities reach a persistent equilibrium (Barton et al. 2013; Butterworth et al. 2022). Heterotrophic succession has been described more specifically for fruitassociated invertebrates, where progressive changes in fruit condition alter opportunities for colonization and resource use (Souza-silva and Ferreira 2004). Previous studies of Psidium guajava and baobab fruits have likewise documented changes in richness, abundance or assemblage composition among fruit conditions or stages (Łukasik and Johnson 2007; Silva and Ferreira 2011). Our results extend this framework by repeatedly following the same standardized fallen-fruit patches from fresh condition through advanced degradation. The resulting trajectory shows that substantial community turnover can occur within only a few weeks as the resource itself changes.

The fresh and pre-rotten stages represented the main period of invertebrate diversity and resource use. Drosophilid and tephritid flies were particularly associated with fresh fruits, whereas several nitidulid beetles and ants were indicators of the combined fresh and pre-rotten stages. At placement, the fruit surface was punctured or broken by impact, exposing pulp and creating immediate access to juice and internal tissues. As decomposition progressed into the pre-rotten stage, softening tissue and the development of rotten odour coincided with continued high richness and peak relative abundance. Earlier studies similarly show that changes in fruit condition can alter the establishment and abundance of fruit-associated invertebrates (Silva and Ferreira 2011; Souza-silva and Ferreira 2004), while fallen fruits can remain important resources for fruit-associated insects under particular environmental conditions (Deconninck et al. 2024). Because fruit chemistry, microbial activity and fermentation were not measured directly in our study, however, the mechanisms attracting particular taxa cannot be distinguished. The observed pattern nevertheless indicates that the early and intermediate stages provided the broadest opportunities for exploitation by the assemblage recorded here.

As fruits progressed beyond the pre-rotten stage, declining diversity was accompanied by clear taxonomic replacement. Unidentified Staphylinidae were associated with the pre-rotten and post-rotten stages, Collembola with post-rotten and degraded fruits, and Blattodea with the broader post-fresh period; the abundance-weighted analysis additionally identified Metochus uniguttatus as a degraded-stage indicator. By these later stages, much of the pulp had been lost and the remaining fruit material had become strongly decomposed or dry. Consequently, later fruit-days were not simply depauperate versions of the fresh-fruit assemblage but supported a compositionally different subset of taxa. This is consistent with studies of other heterotrophic or ephemeral systems in which resource transformation through decomposition is associated with temporal replacement of arthropod assemblages (Koskinen et al. 2022; Lettenmaier et al. 2025; Nadeau et al. 2015). In the present system, succession therefore involved two related processes: a progressive reduction in the number and intensity of resource users and a shift in the identity of taxa occupying the fruit patch.

The multivariate analyses reinforce this interpretation, although they also qualify it. Decomposition stage accounted for substantially more variation in both abundance-weighted and presence–absence community composition than did microhabitat. At the same time, multivariate dispersion differed among decomposition stages, with degraded fruit-days showing lower compositional dispersion than earlier stages. The PERMANOVA result should therefore be understood as reflecting both a shift in assemblage composition and a change in within-stage variability (Anderson, Ellingsen, and Mcardle 2006; Anderson and Walsh 2013). Ecologically, the latter pattern is also informative. As the fruit resource approached depletion, fewer taxa remained and degraded fruit-days occupied a more restricted region of community space (Anderson et al. 2006). The agreement among the abundance-weighted analysis, presence–absence analysis and indicator-taxon results nevertheless provides convergent evidence that community structure changed systematically during decomposition (Anderson et al. 2006; Barton et al. 2013; Lettenmaier et al. 2025).

Against this strong temporal signal, microhabitat explained comparatively little variation. Exposed and shaded fruits showed some differences in community composition, but these were much smaller than differences associated with decomposition stage, while direct microhabitat effects on richness and total peak relative abundance were weak. Environmental conditions can modify successional trajectories on ephemeral resources; for example, experimental work on dead wood has shown that environmental context can alter beetle succession during resource decay (Vindstad et al. 2020). In our study, exposed and shaded sites could plausibly have differed in heating, moisture retention or desiccation, but these mechanisms were not measured directly at the fruit level. The observed differences may also have arisen from variation in colonization, local ant activity or other biotic conditions. Moreover, microhabitat was represented by only four fruits in each category. The exposed–shaded contrast should therefore be interpreted as a secondary microhabitat-associated pattern, rather than as evidence that shade, temperature or moisture directly drove assemblage differences (Bracewell, Johnston, and Clark 2024; Kim et al. 2021; Sbaraglia et al. 2025). What is clear from the present data is that the temporal transformation of the fruit resource had a much stronger association with assemblage structure than this broad spatial contrast.

Ants formed a conspicuous component of these temporary resource patches and added a behavioural dimension to the successional pattern. Camponotus crassisquamis was observed carrying pulp fragments containing seeds during early decomposition, while Pheidole sp. 1 exploited fruits across early and intermediate stages and was itself an indicator of the combined fresh and pre-rotten stages. Pheidole workers fed on exposed pulp, deposited soil or leaf fragments over parts of the fruit, guarded access points and attacked or carried some larvae and coleopterans. Ants are major agents of resource removal in tropical systems (Agaldo, Christianini, and Chapman 2021; Griffiths et al. 2017), and their interactions with concentrated organic resources can extend beyond feeding alone (King et al. 2018; Neumann and Pinter-Wollman 2022). Whether these conspicuous behaviours translated into measurable effects on the wider fruit assemblage, however, was less clear. After decomposition stage and other covariates were accounted for, associations between Pheidole sp. 1 peak relative abundance and non-ant richness or peak relative abundance were weak or marginal, and taxon-specific models provided no consistent evidence of negative associations (Costa-Silva, Moreira, and Del-Claro 2023; Ruzi et al. 2021). The behavioural observations therefore suggest that Pheidole can physically modify and defend fruit patches, but the present data do not demonstrate community-level suppression. In a rapidly changing resource system, both ants and other taxa may independently track the same underlying decomposition gradient, making causal interactions difficult to infer from co-occurrence patterns alone (Estrepo and Omez 1999; Lima, Oliveira, and Silveira 2013; Nooten et al. 2022; Passos and Oliveira 2003). Experimental ant exclusion or controlled-access experiments would be needed to determine whether Pheidole directly alters colonization, persistence or resource use by other invertebrates.

Beyond their possible interactions with other invertebrates, ants also altered the physical fate of fruit material by removing pulp and seeds. Recorded removal distances ranged from 0.035 to 8.37 m, with the longest movement attributed to Pheidole sp. 1. Such observations fit a broader literature showing that ants interact with and remove diaspores of plants lacking specialized adaptations for ant dispersal (Anjos et al. 2020; Lima et al. 2013). A meta-analysis further demonstrates that ants can contribute substantially to the secondary removal of non-myrmecochorous diaspores, although removal distances and outcomes vary among systems (Anjos et al. 2020). Our observations should nevertheless be interpreted specifically as pulp or seed removal rather than confirmed seed dispersal. Seeds were not followed after transport, and we did not measure deposition site quality, germination or seedling establishment. Whether ant-mediated removal benefits, harms or has no effect on P. guajava recruitment therefore remains unknown. The observations instead identify a natural-history pathway through which consumers exploiting a decomposing fruit can redistribute components of the resource beyond the original patch.

Several limitations define the scope of these conclusions. Most importantly, the experiment comprised eight fruits within a single tropical grove. Repeated observations generated 141 fruit-days and provided fine temporal resolution of each fruit’s decomposition trajectory, but these fruit-days are not equivalent to 141 independent spatial replicates. Generalization beyond this site, fruit species or set of experimental patches should therefore remain cautious. Second, abundance was represented by the highest number of individuals observed during hourly scans. This peak relative abundance index provides a standardized measure of maximum observed daily resource use, but it is not an estimate of total daily abundance and may underrepresent short visits or activity outside daylight observation periods (Burton et al. 2015; Siegel and Dee 2025; Steenweg et al. 2017). Third, microhabitat was classified broadly rather than characterized through continuous measurements of fruit temperature, moisture, canopy openness or microbial succession. Several taxa also remained identified as morphospecies or higher taxonomic units, limiting species-specific ecological interpretation. Finally, the ant analyses were observational and the seed-removal observations opportunistic; neither can establish causal effects (Arif and Macneil 2022; Byrnes and Dee 2024). These limitations constrain mechanistic interpretation, but they do not change the central observation that assemblage structure shifted strongly and repeatedly as individual fruits passed through successive decomposition stages.

Overall, this study shows that a single fallen fruit can pass through several biologically distinct resource states within only a few weeks. Fresh and pre-rotten guavas supported diverse and intensively used assemblages dominated by fruit-associated flies, beetles and ants, whereas later decomposition stages supported fewer but compositionally different taxa. The resulting pattern is consistent with heterotrophic succession (Seibold et al. 2022; Souza-Silva and Ferreira 2004) driven primarily by temporal change in the resource itself, with microhabitat providing a weaker secondary context and ants contributing additional resource modification and removal. More broadly, fallen fruits should not be viewed simply as discarded plant material: they constitute small, transient patches that add temporal as well as spatial heterogeneity to tropical invertebrate communities (Butterworth et al. 2022; Deconninck et al. 2024). Their short lifespan and rapid biological turnover make them particularly tractable systems (Ferrari et al. 2017; Vindstad et al. 2020) for examining colonization, resource depletion and community replacement at fine temporal scales.

## Acknowledgements

We thank the authorities and residents of Barasat Kisoloy for permitting and supporting the fieldwork, and the fruit vendors who supplied fruit for the study. We are grateful to the Director of the Zoological Survey of India for permitting us to consult the institute’s scientists for species identification, and to Dr. S. Sheela for her assistance with sorting specimens and arranging their identification. We thank Dr. Kuldeep Roy for his help during the study and Dr. Udipta Chakraborty for his comments that improved the manuscript. This paper was typeset with the bioRxiv word template by @Chrelli: www.github.com/chrelli/bioRxiv-word-template

## Author contributions

Silanjan Bhattacharya conceptualised and supervised the study. Sourabh Biswas conceptualised, developed the methodology, conducted the study, curated and analysed the data, and drafted the manuscript. Silanjan Bhattacharya approved the final manuscript.

## Ethics approval and permits

Fieldwork was conducted at Barasat Kisoloy, West Bengal, in 2016 with the permission of the site authority. Invertebrates were primarily identified through field observations and photographs. Specimens were collected only when identification by those methods was insufficient and were submitted to the Zoological Survey of India, Kolkata, for expert identification. No vertebrates or human participants were involved.

## Data and code availability statement

The data supporting this study will be archived in a public repository. The repository link and DOI will be added when the deposit is complete.

## Supplementary materials

**Table S1.**
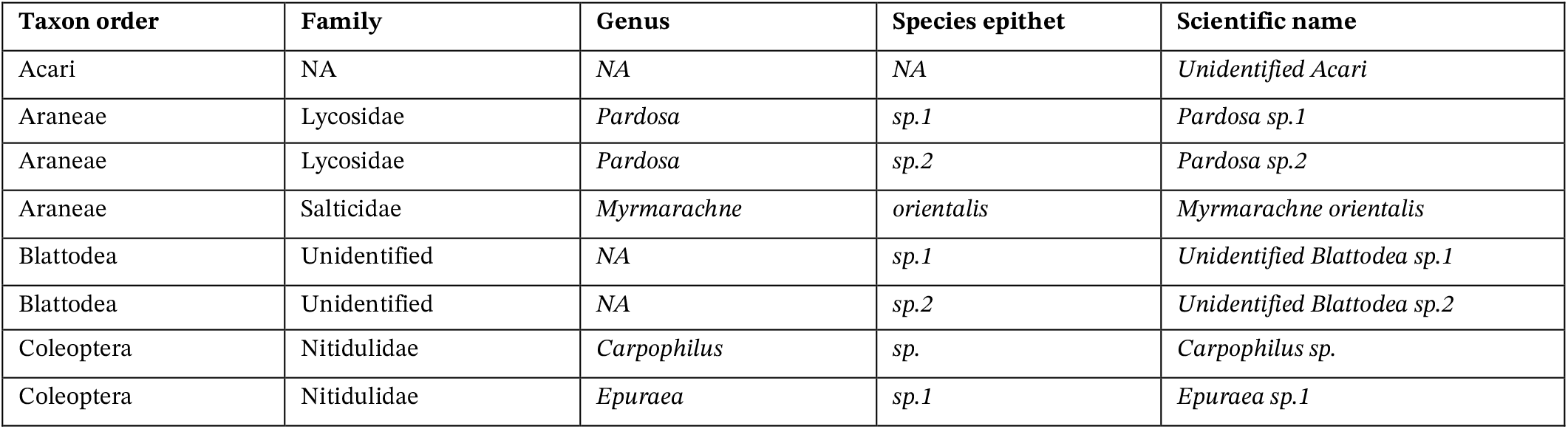
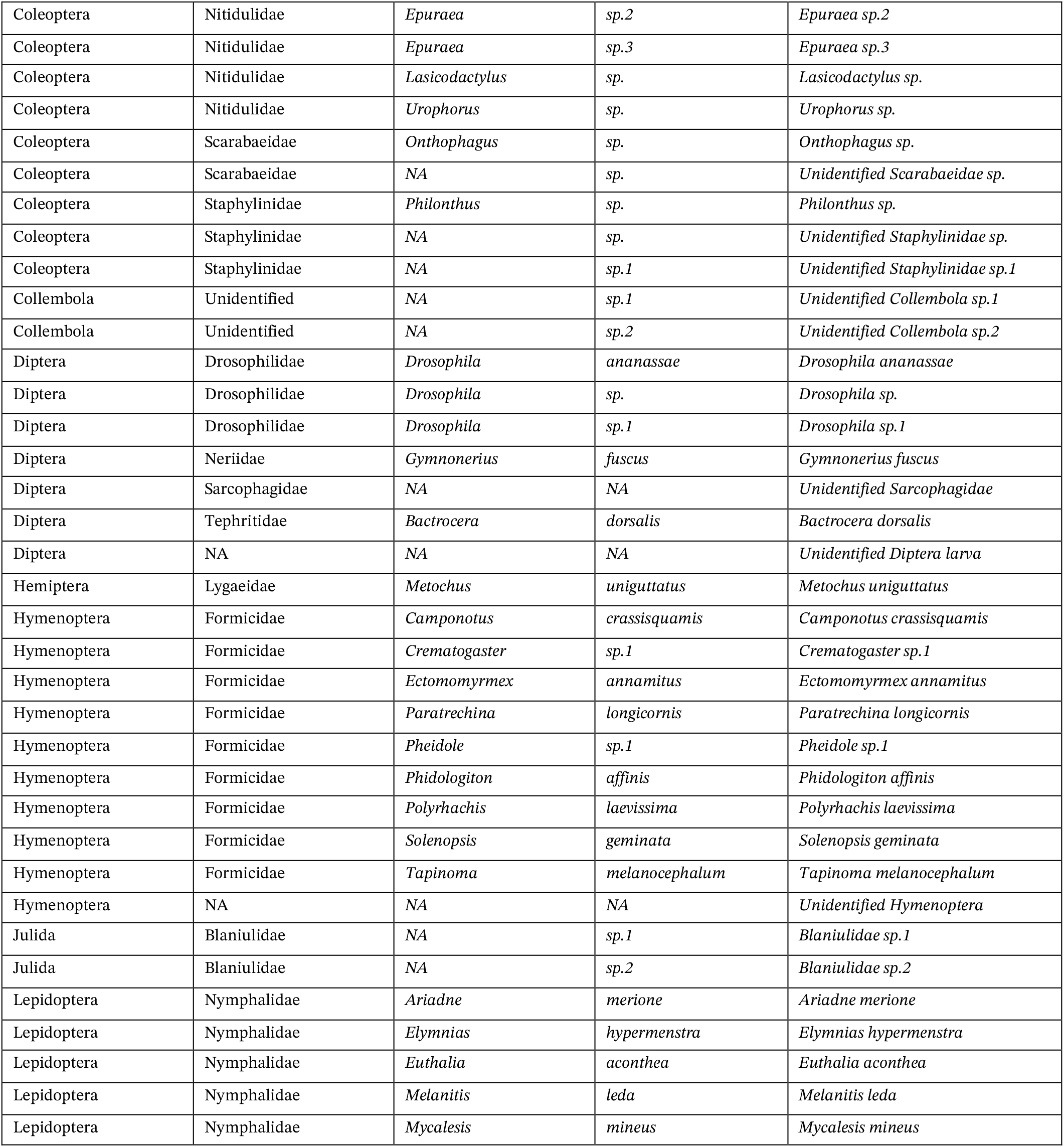
Full taxon list recorded from fallen ripe guava fruits. List of all 44 invertebrate taxa recorded during the decomposition of fallen ripe *Psidium guajava* fruits in exposed and shaded microhabitats. For each taxon, the table provides taxonomic order, family, genus, species epithet or morphospecies name. Taxa not identified to species level were retained as morphospecies or higher taxonomic units and used consistently across analyses.

| Taxon order | Family | Genus | Species epithet | Scientific name |
| --- | --- | --- | --- | --- |
| Acari | NA | NA | NA | Unidentified Acari |
| Araneae | Lycosidae | *Pardosa* | *sp.1* | *Pardosa sp.1* |
| Araneae | Lycosidae | *Pardosa* | *sp.2* | *Pardosa sp.2* |
| Araneae | Salticidae | *Myrmarachne* | *orientalis* | *Myrmarachne orientalis* |
| Blattodea | Unidentified | NA | *sp.1* | Unidentified Blattodea sp.1 |
| Blattodea | Unidentified | NA | *sp.2* | Unidentified Blattodea sp.2 |
| Coleoptera | Nitidulidae | *Carpophilus* | *sp.* | *Carpophilus sp.* |
| Coleoptera | Nitidulidae | *Epuraea* | *sp.1* | *Epuraea sp.1* |
| Coleoptera | Nitidulidae | *Epuraea* | *sp.2* | *Epuraea sp.2* |
| Coleoptera | Nitidulidae | *Epuraea* | *sp.3* | *Epuraea sp.3* |
| Coleoptera | Nitidulidae | *Lasicodactylus* | *sp.* | *Lasicodactylus sp.* |
| Coleoptera | Nitidulidae | *Urophorus* | *sp.* | *Urophorus sp.* |
| Coleoptera | Scarabaeidae | *Onthophagus* | *sp.* | *Onthophagus sp.* |
| Coleoptera | Scarabaeidae | NA | *sp.* | Unidentified Scarabaeidae *sp.* |
| Coleoptera | Staphylinidae | *Philonthus* | *sp.* | *Philonthus sp.* |
| Coleoptera | Staphylinidae | NA | *sp.* | Unidentified Staphylinidae *sp.* |
| Coleoptera | Staphylinidae | NA | *sp.1* | Unidentified Staphylinidae *sp.1* |
| Collembola | Unidentified | NA | *sp.1* | Unidentified Collembola *sp.1* |
| Collembola | Unidentified | NA | *sp.2* | Unidentified Collembola *sp.2* |
| Diptera | Drosophilidae | *Drosophila* | *ananassae* | *Drosophila ananassae* |
| Diptera | Drosophilidae | *Drosophila* | *sp.* | *Drosophila sp.* |
| Diptera | Drosophilidae | *Drosophila* | *sp.1* | *Drosophila sp.1* |
| Diptera | Neriidae | *Gymnonerius* | *fuscus* | *Gymnonerius fuscus* |
| Diptera | Sarcophagidae | NA | NA | Unidentified Sarcophagidae |
| Diptera | Tephritidae | *Bactrocera* | *dorsalis* | *Bactrocera dorsalis* |
| Diptera | NA | NA | NA | Unidentified Diptera larva |
| Hemiptera | Lygaeidae | *Metochus* | *uniguttatus* | *Metochus uniguttatus* |
| Hymenoptera | Formicidae | *Camponotus* | *crassisquamis* | *Camponotus crassisquamis* |
| Hymenoptera | Formicidae | *Crematogaster* | *sp.1* | *Crematogaster sp.1* |
| Hymenoptera | Formicidae | *Ectomomyrmex* | *annamitus* | *Ectomomyrmex annamitus* |
| Hymenoptera | Formicidae | *Paratrechina* | *longicornis* | *Paratrechina longicornis* |
| Hymenoptera | Formicidae | *Pheidole* | *sp.1* | *Pheidole sp.1* |
| Hymenoptera | Formicidae | *Phidologiton* | *affinis* | *Phidologiton affinis* |
| Hymenoptera | Formicidae | *Polyrhachis* | *laevissima* | *Polyrhachis laevissima* |
| Hymenoptera | Formicidae | *Solenopsis* | *geminata* | *Solenopsis geminata* |
| Hymenoptera | Formicidae | *Tapinoma* | *melanocephalum* | *Tapinoma melanocephalum* |
| Hymenoptera | NA | NA | NA | Unidentified Hymenoptera |
| Julida | Blaniulidae | NA | *sp.1* | *Blaniulidae sp.1* |
| Julida | Blaniulidae | NA | *sp.2* | *Blaniulidae sp.2* |
| Lepidoptera | Nymphalidae | *Ariadne* | *merione* | *Ariadne merione* |
| Lepidoptera | Nymphalidae | *Elymnias* | *hypermenstra* | *Elymnias hypermenstra* |
| Lepidoptera | Nymphalidae | *Euthalia* | *aconthea* | *Euthalia aconthea* |
| Lepidoptera | Nymphalidae | *Melanitis* | *leda* | *Melanitis leda* |
| Lepidoptera | Nymphalidae | *Mycalesis* | *mineus* | *Mycalesis mineus* |

**Table S2.**
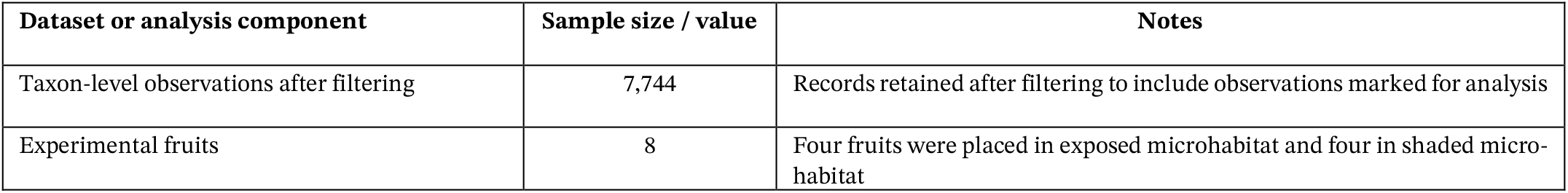
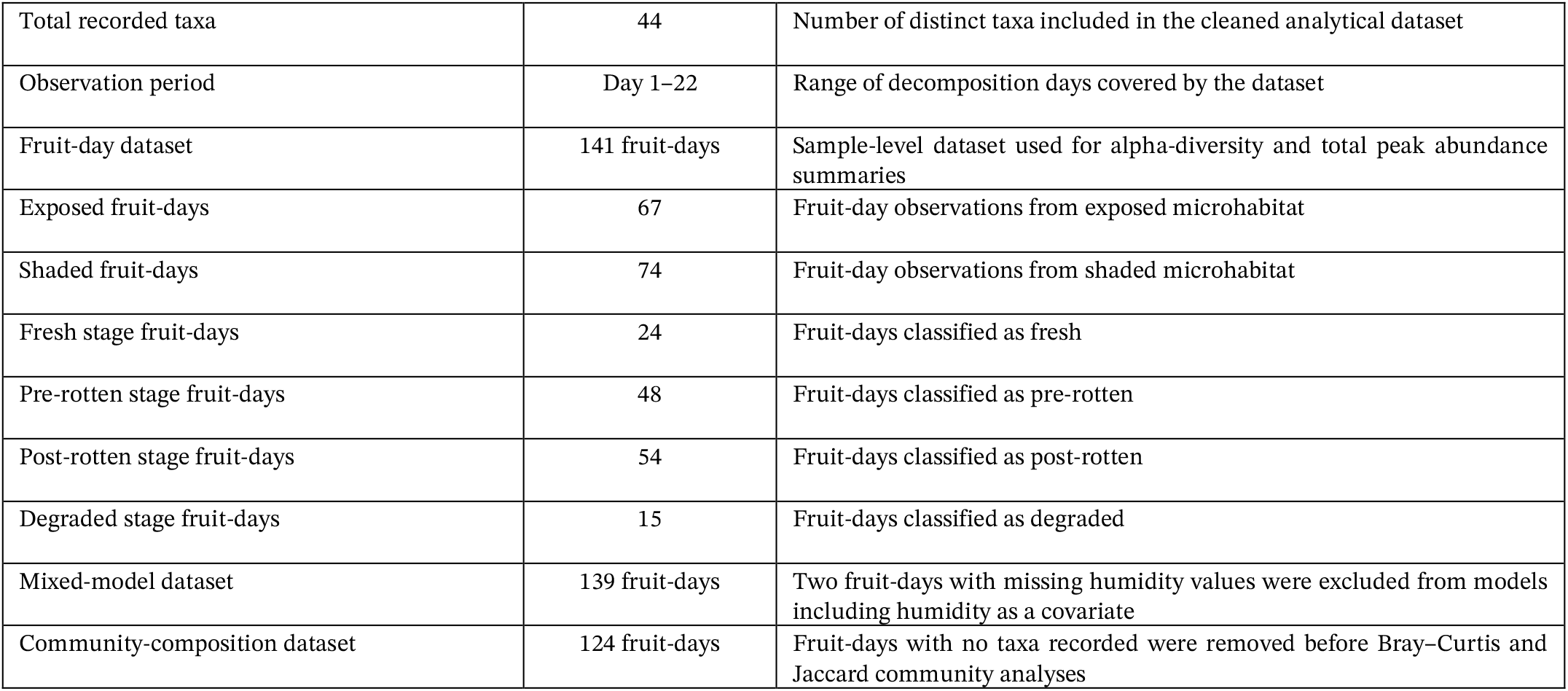
Dataset structure and analysis-specific sample sizes used in the guava fallen-fruit heterotrophic succession study. Taxon-level observations were first filtered to retain records marked for analysis and were then summarized at the fruit-day level. Alpha-diversity and total peak relative abundance analyses used the fruit-day dataset, whereas mixed models excluded two fruit-days with missing humidity values. Community-composition analyses excluded fruit-days with no taxa recorded before calculating Bray–Curtis and Jaccard dissimilarities.

| Dataset or analysis component | Sample size / value | Notes |
| --- | --- | --- |
| Taxon-level observations after filtering | 7,744 | Records retained after filtering to include observations marked for analysis |
| Experimental fruits | 8 | Four fruits were placed in exposed microhabitat and four in shaded micro-habitat |
| Total recorded taxa | 44 | Number of distinct taxa included in the cleaned analytical dataset |
| Observation period | Day 1–22 | Range of decomposition days covered by the dataset |
| Fruit-day dataset | 141 fruit-days | Sample-level dataset used for alpha-diversity and total peak abundance summaries |
| Exposed fruit-days | 67 | Fruit-day observations from exposed microhabitat |
| Shaded fruit-days | 74 | Fruit-day observations from shaded microhabitat |
| Fresh stage fruit-days | 24 | Fruit-days classified as fresh |
| Pre-rotten stage fruit-days | 48 | Fruit-days classified as pre-rotten |
| Post-rotten stage fruit-days | 54 | Fruit-days classified as post-rotten |
| Degraded stage fruit-days | 15 | Fruit-days classified as degraded |
| Mixed-model dataset | 139 fruit-days | Two fruit-days with missing humidity values were excluded from models including humidity as a covariate |
| Community-composition dataset | 124 fruit-days | Fruit-days with no taxa recorded were removed before Bray–Curtis and Jaccard community analyses |

**Table S3A.** Fixed-effect estimates from the accepted negative-binomial GLMM for taxon richness. Table S3.Fixed-effect estimates from the accepted negative-binomial generalized linear mixed model for taxon richness per fruit-day. The model used a negative-binomial error distribution with a log link. Exposed microhabitat and fresh decomposition stage were used as reference categories. Fruit identity was included as a random intercept to account for repeated sampling of the same fruits through decomposition. Positive estimates indicate higher expected taxon richness relative to the reference level on the log scale, whereas negative estimates indicate lower expected richness. **Model formula:** Taxon richness ~ microhabitat × decomposition stage + standardized temperature + standardized humidity + standardized fruit weight + (1 | fruit identity)

| Predictor | Estimate | Std. Error | z value | p-value |
| --- | --- | --- | --- | --- |
| Intercept | 2.33200 | 0.22477 | 10.375 | < 2e-16 |
| Microhabitat: Shaded | -0.29340 | 0.30405 | -0.965 | 0.334558 |
| Decomposition stage: Pre-rotten | -0.01043 | 0.17097 | -0.061 | 0.951345 |
| Decomposition stage: Post-rotten | -0.66433 | 0.19140 | -3.471 | 0.000519 |
| Decomposition stage: Degraded | -1.65492 | 0.31153 | -5.312 | 1.08e-07 |
| Temperature, standardized | -0.07132 | 0.10707 | -0.666 | 0.505347 |
| Humidity, standardized | -0.07970 | 0.06540 | -1.219 | 0.222956 |
| Fruit weight, standardized | -0.19894 | 0.14355 | -1.386 | 0.165783 |
| Shaded × Pre-rotten | 0.18708 | 0.22409 | 0.835 | 0.403822 |
| Shaded × Post-rotten | 0.24043 | 0.23738 | 1.013 | 0.311139 |
| Shaded × Degraded | -0.52090 | 0.48943 | -1.064 | 0.287191 |

**Table S3B.** Model-estimated taxon richness across decomposition stages of fallen guava fruits.

| Decomposition stage | Estimated richness | SE | 95% lower CI | 95% upper CI |
| --- | --- | --- | --- | --- |
| Fresh | 8.87 | 1.5 | 6.37 | 12.36 |
| Pre-rotten | 9.64 | 1.34 | 7.35 | 12.65 |
| Post-rotten | 5.15 | 0.75 | 3.88 | 6.84 |
| Degraded | 1.31 | 0.34 | 0.78 | 2.19 |

**Table S3C.**
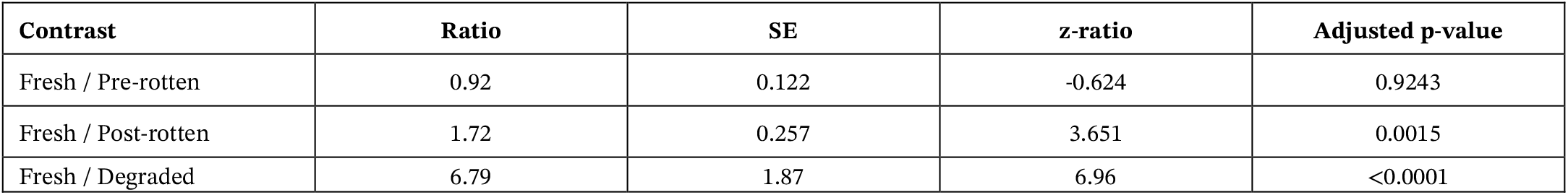
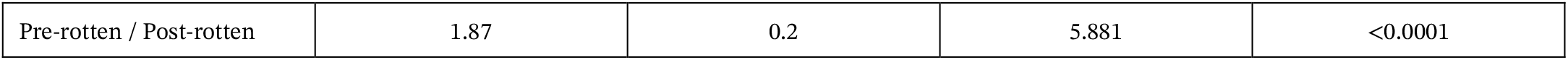
Tukey-adjusted pairwise comparisons of model-estimated richness among decomposition stages.

| Contrast | Ratio | SE | z-ratio | Adjusted p-value |
| --- | --- | --- | --- | --- |
| Fresh / Pre-rotten | 0.92 | 0.122 | -0.624 | 0.9243 |
| Fresh / Post-rotten | 1.72 | 0.257 | 3.651 | 0.0015 |
| Fresh / Degraded | 6.79 | 1.87 | 6.96 | <0.0001 |
| Pre-rotten / Post-rotten | 1.87 | 0.2 | 5.881 | <0.0001 |

**Table S4A.** Fixed-effect estimates from the accepted linear mixed model for Shannon diversity. Fixed-effect estimates and model summary from the accepted linear mixed model for Shannon diversity per fruit-day. The model was fitted using maximum likelihood. Exposed microhabitat and fresh decomposition stage were used as reference categories. Fruit identity was included as a random intercept to account for repeated sampling of the same fruits through decomposition. Negative estimates indicate lower Shannon diversity relative to the reference level, whereas positive estimates indicate higher Shannon diversity. Model formula: Shannon diversity ~ microhabitat × decomposition stage + standardized temperature + standardized humidity + standardized fruit weight + (1 | fruit identity)

| Predictor | Estimate | Std. Error | t value |
| --- | --- | --- | --- |
| Intercept | 1.45847 | 0.18696 | 7.801 |
| Microhabitat: Shaded | 0.03578 | 0.24424 | 0.147 |
| Decomposition stage: Pre-rotten | -0.22943 | 0.19783 | -1.160 |
| Decomposition stage: Post-rotten | -0.63455 | 0.20408 | -3.109 |
| Decomposition stage: Degraded | -1.40724 | 0.28379 | -4.959 |
| Temperature, standardized | -0.03885 | 0.10232 | -0.380 |
| Humidity, standardized | -0.10444 | 0.06736 | -1.550 |
| Fruit weight, standardized | -0.05490 | 0.09625 | -0.570 |
| Shaded × Pre-rotten | 0.06510 | 0.25889 | 0.251 |
| Shaded × Post-rotten | 0.22856 | 0.25563 | 0.894 |
| Shaded × Degraded | 0.04223 | 0.35353 | 0.119 |

**Table S4B.** Analysis of variance table for the Shannon diversity mixed model.

| Fixed effect | npar | Sum Sq | Mean Sq | F value |
| --- | --- | --- | --- | --- |
| Microhabitat | 1 | 0.0993 | 0.0993 | 0.3731 |
| Decomposition stage | 3 | 22.0236 | 7.3412 | 27.5726 |
| Temperature | 1 | 0.2337 | 0.2337 | 0.8777 |
| Humidity | 1 | 0.6237 | 0.6237 | 2.3425 |

**Table S4C.** Model-estimated Shannon diversity by microhabitat and decomposition stage.

| Microhabitat | Decomposition stage | Estimated Shannon diversity | SE | df | 95% lower CI | 95% upper CI |
| --- | --- | --- | --- | --- | --- | --- |
| Exposed | Fresh | 1.458 | 0.214 | 81.5 | 1.033 | 1.883 |
| Shaded | Fresh | 1.4938 | 0.214 | 84.8 | 1.067 | 1.92 |
| Exposed | Pre-rotten | 1.2286 | 0.168 | 34.4 | 0.887 | 1.57 |
| Shaded | Pre-rotten | 1.3295 | 0.168 | 35.8 | 0.989 | 1.67 |
| Exposed | Post-rotten | 0.8235 | 0.169 | 36 | 0.48 | 1.166 |
| Shaded | Post-rotten | 1.0878 | 0.163 | 31.4 | 0.756 | 1.42 |
| Exposed | Degraded | 0.0508 | 0.264 | 121 | -0.472 | 0.574 |
| Shaded | Degraded | 0.1288 | 0.231 | 98.2 | -0.33 | 0.587 |

**Table S5.** Fixed-effect estimates from the accepted COM-Poisson GLMM for total peak relative abundance index. Fixed-effect estimates and model summary from the accepted COM-Poisson generalized linear mixed model for total peak relative abundance index per fruit-day. The model used a COM-Poisson error distribution with a log link. Exposed microhabitat and fresh decomposition stage were used as reference categories. Fruit identity was included as a random intercept to account for repeated sampling of the same fruits through decomposition. Positive estimates indicate higher expected total peak relative abundance relative to the reference level on the log scale, whereas negative estimates indicate lower expected total peak relative abundance. **Model formula:** Total peak relative abundance index ~ microhabitat × decomposition stage + standardized temperature + standardized humidity + standardized fruit weight + (1 | fruit identity)

| Predictor | Estimate | Std. Error | z value | p-value |
| --- | --- | --- | --- | --- |
| Intercept | 4.9473 | 0.2594 | 19.074 | < 2e-16 |
| Microhabitat: Shaded | -0.4849 | 0.3570 | -1.358 | 0.1743 |
| Decomposition stage: Pre-rotten | -0.1662 | 0.2420 | -0.687 | 0.4923 |
| Decomposition stage: Post-rotten | -1.8964 | 0.2908 | -6.522 | 6.94e-11 |
| Decomposition stage: Degraded | -3.0743 | 0.4862 | -6.323 | 2.57e-10 |
| Temperature, standardized | 0.1165 | 0.1536 | 0.758 | 0.4484 |
| Humidity, standardized | 0.1303 | 0.0994 | 1.311 | 0.1900 |
| Fruit weight, standardized | -0.0723 | 0.1591 | -0.454 | 0.6495 |
| Shaded × Pre-rotten | 0.5576 | 0.3293 | 1.693 | 0.0904 |
| Shaded × Post-rotten | 0.8395 | 0.3754 | 2.237 | 0.0253 |
| Shaded × Degraded | -0.3081 | 0.6437 | -0.479 | 0.6322 |

**Figure S2.**
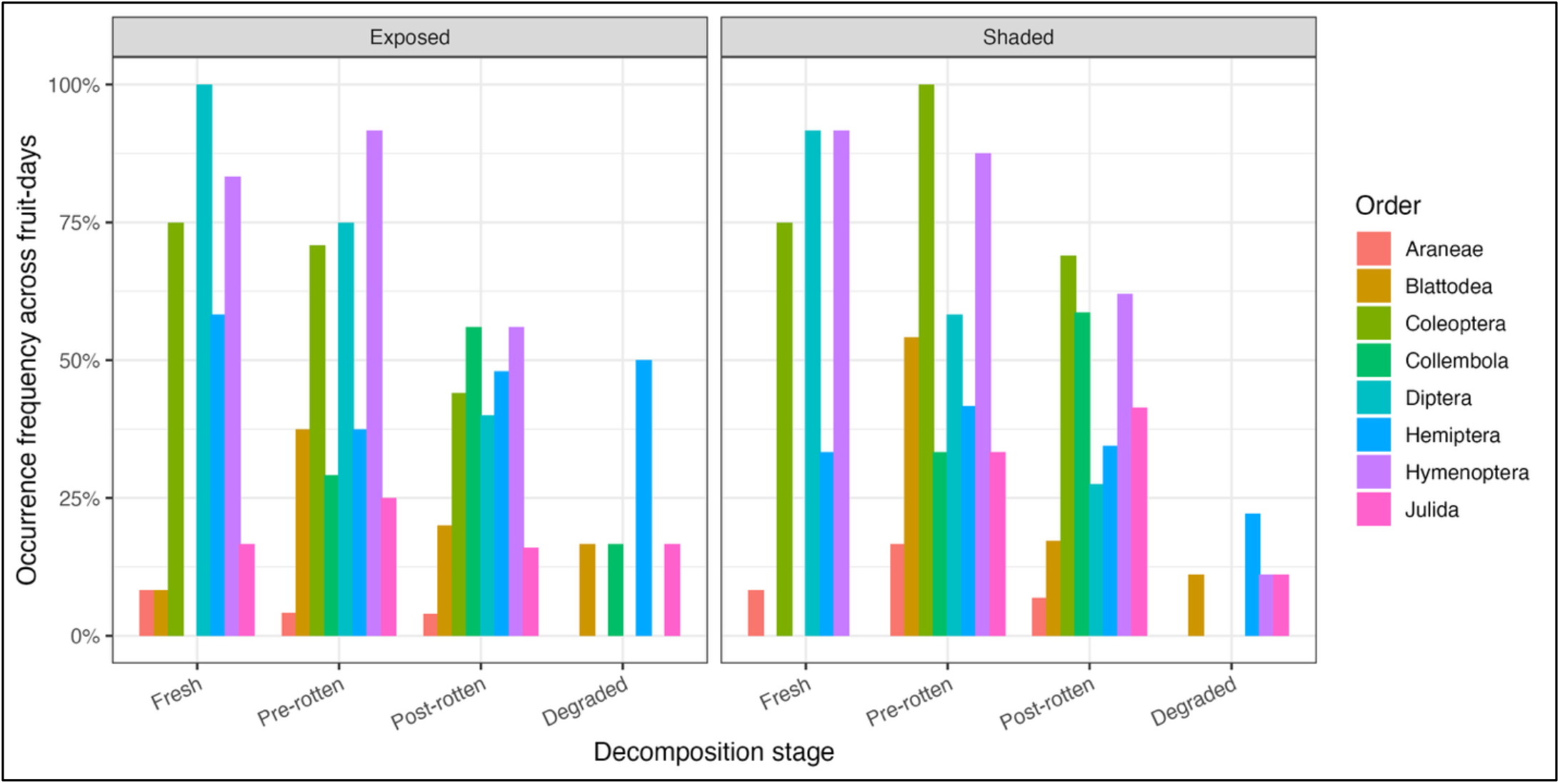
Order-level occurrence frequency across decomposition stages and microhabitats. Occurrence frequency of major invertebrate orders recorded on fallen ripe Psidium guajava fruits across fruit-days in exposed and shaded microhabitats. Bars show the percentage of fruit-days within each decomposition stage and microhabitat in which at least one individual of a given order was recorded. Early decomposition stages were characterized by frequent occurrence of Diptera, Coleoptera and Hymenoptera, whereas later post-rotten and degraded stages showed lower overall occurrence and increased representation of groups such as Hemiptera, Collembola, Blattodea and Julida.

**Figure S3.**
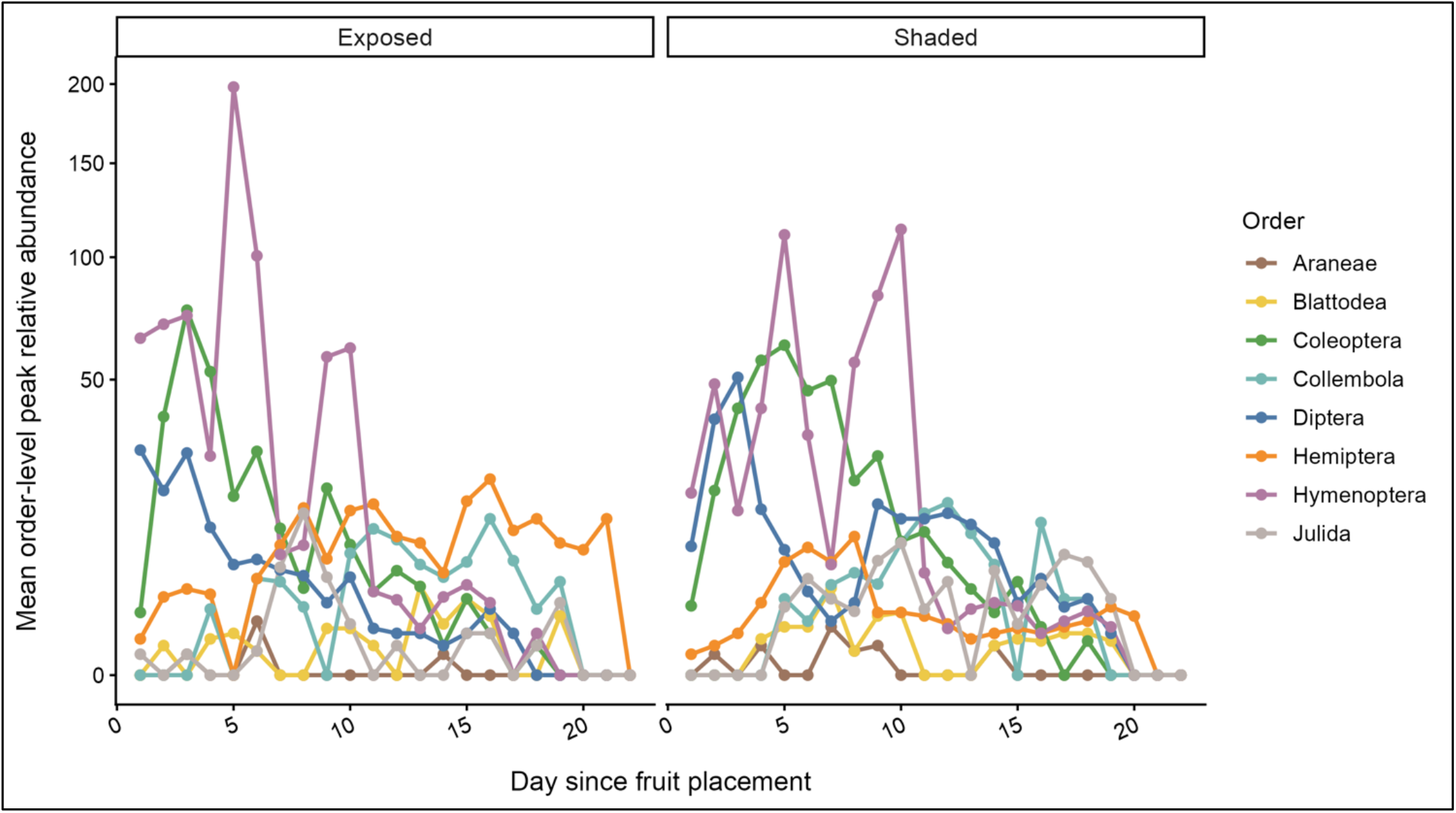
Day-wise order-level succession on fallen ripe guava fruits. Mean order-level peak relative abundance of major invertebrate orders recorded on fallen ripe Psidium guajava fruits across days since fruit placement in exposed and shaded microhabitats. Points and lines show the mean daily peak relative abundance index for each order, calculated from scan-based maximum daily counts. Early decomposition days were characterized by high activity of Hymenoptera, Coleoptera and Diptera, whereas later days showed lower and more variable activity, with increased contributions from Hemiptera, Collembola, Blattodea and Julida in some periods. Because values represent peak relative abundance rather than total abundance, the figure should be interpreted as a descriptive visualization of temporal order-level succession.

**Table S6A.**
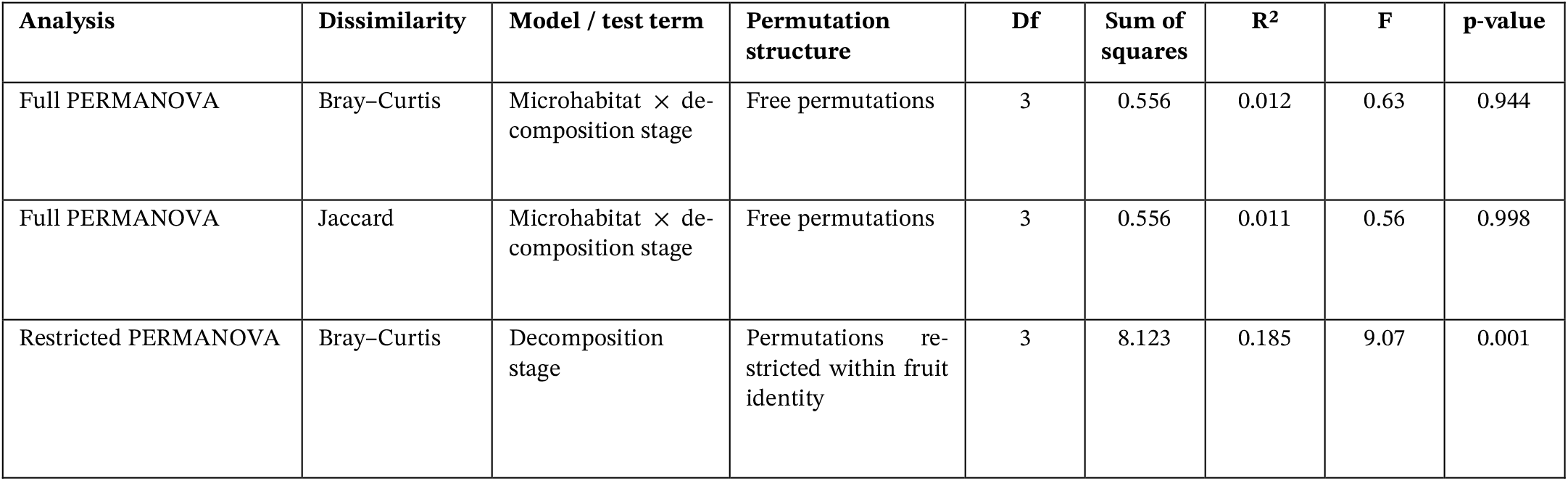
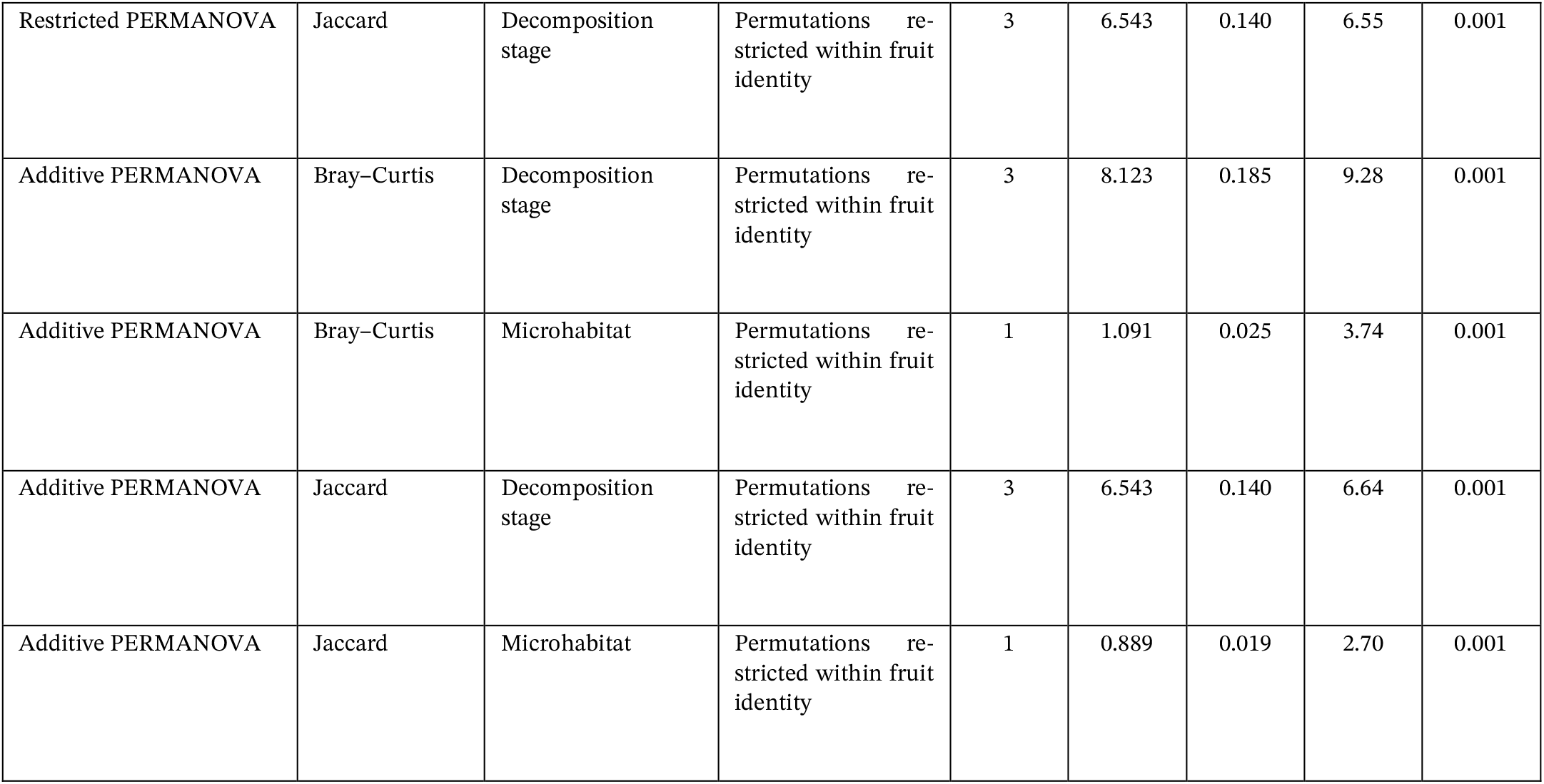
PERMANOVA and multivariate dispersion results for community-composition analyses. PERMANOVA and multivariate dispersion results for fruit-associated invertebrate community composition across decomposition stages and microhabitats. Bray–Curtis dissimilarity was calculated from square-root-transformed peak relative abundance, whereas Jaccard dissimilarity was calculated from presence/absence data. Full interaction PERMANOVA models tested whether the compositional trajectory across decomposition stages differed between exposed and shaded microhabitats. Restricted stage-only and additive PERMANOVA models used permutations constrained within fruit identity to account for repeated sampling of the same fruits through decomposition. Homogeneity of multivariate dispersion was tested using Bray–Curtis distances. Decomposition stages differed significantly in dispersion, whereas exposed and shaded microhabitats did not. Pairwise dispersion comparisons are Tukey-adjusted. **PERMANOVA model formulas:** **Full interaction model:** community dissimilarity ~ microhabitat × decomposition stage **Restricted stage-only model:** community dissimilarity ~ decomposition stage **Additive model:** community dissimilarity ~ decomposition stage + microhabitat

| Analysis | Dissimilarity | Model / test term | Permutation structure | Df | Sum of squares | R<sup>2</sup> | F | p-value |
| --- | --- | --- | --- | --- | --- | --- | --- | --- |
| Full PERMANOVA | Bray–Curtis | Microhabitat × decomposition stage | Free permutations | 3 | 0.556 | 0.012 | 0.63 | 0.944 |
| Full PERMANOVA | Jaccard | Microhabitat × decomposition stage | Free permutations | 3 | 0.556 | 0.011 | 0.56 | 0.998 |
| Restricted PERMANOVA | Bray–Curtis | Decomposition stage | Permutations restricted within fruit identity | 3 | 8.123 | 0.185 | 9.07 | 0.001 |
| Restricted PERMANOVA | Jaccard | Decomposition stage | Permutations restricted within fruit identity | 3 | 6.543 | 0.140 | 6.55 | 0.001 |
| Additive PERMANOVA | Bray–Curtis | Decomposition stage | Permutations restricted within fruit identity | 3 | 8.123 | 0.185 | 9.28 | 0.001 |
| Additive PERMANOVA | Bray–Curtis | Microhabitat | Permutations restricted within fruit identity | 1 | 1.091 | 0.025 | 3.74 | 0.001 |
| Additive PERMANOVA | Jaccard | Decomposition stage | Permutations restricted within fruit identity | 3 | 6.543 | 0.140 | 6.64 | 0.001 |
| Additive PERMANOVA | Jaccard | Microhabitat | Permutations restricted within fruit identity | 1 | 0.889 | 0.019 | 2.70 | 0.001 |

**Table S6B.**
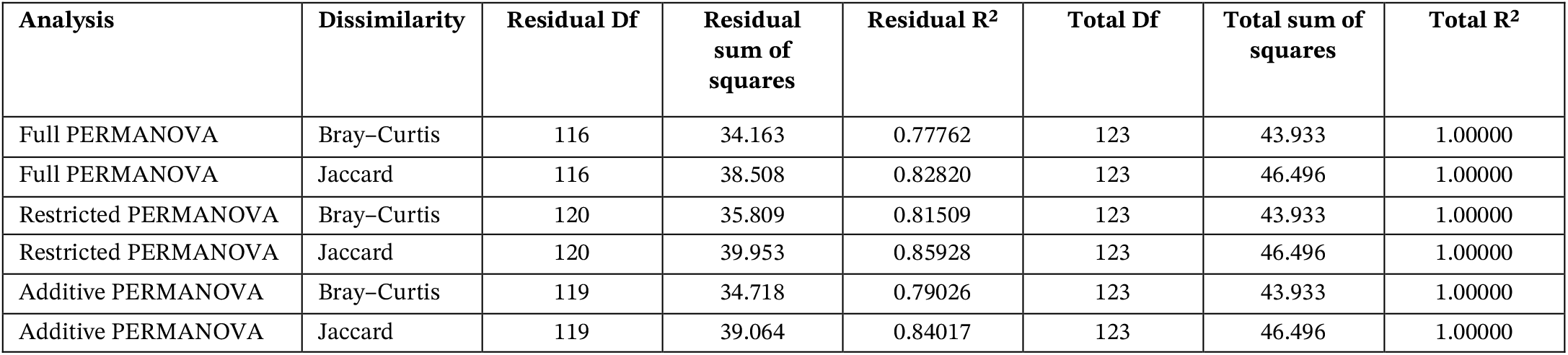
Model residual and total information:

**Table S6C.** Multivariate dispersion tests:

| Dispersion test | Dissimilarity | Grouping variable | Df | Sum of squares | Mean square | F | p-value |
| --- | --- | --- | --- | --- | --- | --- | --- |
| Betadisper ANOVA | Bray–Curtis | Decomposition stage | 3 | 0.42065 | 0.140216 | 12.54 | 3.457e-07 |
| Betadisper permutation test | Bray–Curtis | Decomposition stage | 3 | 0.42065 | 0.140216 | 12.54 | 0.001 |
| Betadisper ANOVA | Bray–Curtis | Microhabitat | 1 | 0.00024 | 0.0002404 | 0.056 | 0.8129 |
| Betadisper permutation test | Bray–Curtis | Microhabitat | 1 | 0.00024 | 0.0002404 | 0.056 | 0.811 |

**Table S6D.**
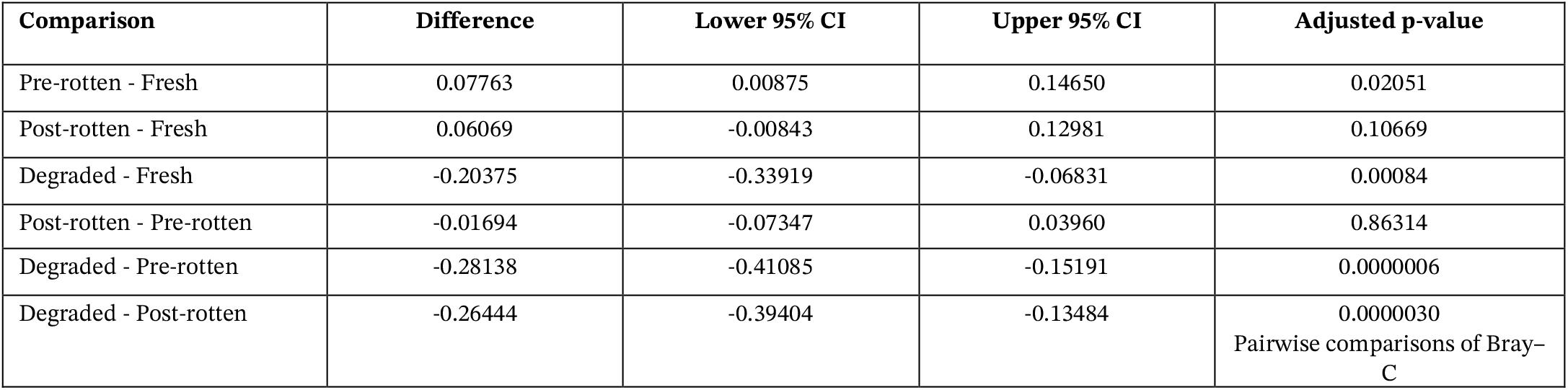
Pairwise comparisons of Bray–Curtis dispersion among decomposition stages:

**Figure S4.**
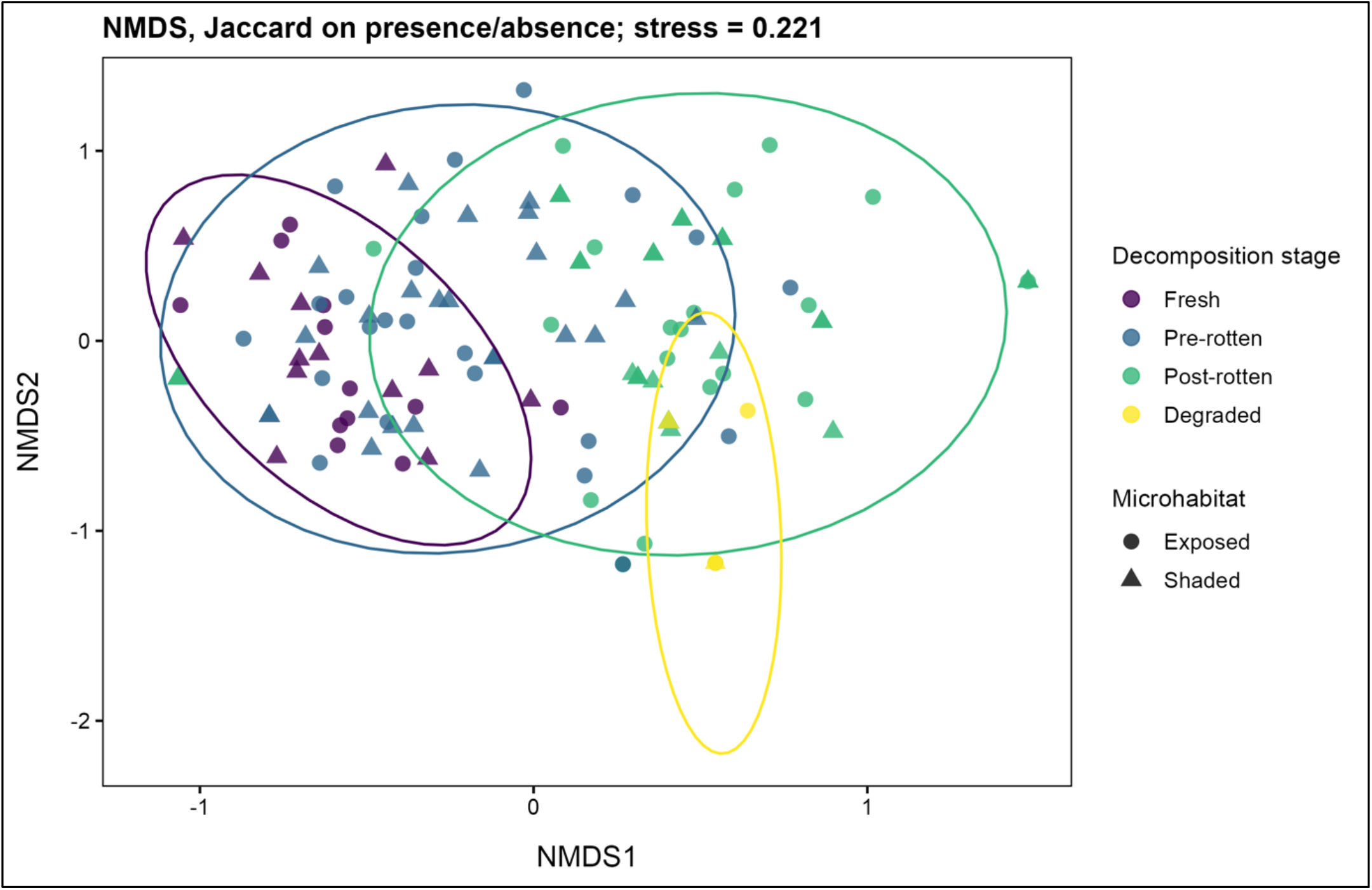
Presence/absence-based community composition of invertebrates across guava fruit decomposition stages. Non-metric multidimensional scaling (NMDS) ordination based on Jaccard dissimilarity of invertebrate presence/absence data from fallen ripe Psidium guajava fruit-days. Points represent individual fruit-days, colours indicate decomposition stage, and shapes indicate microhabitat type: exposed or shaded. Ellipses summarize the dispersion of fruit-days within each decomposition stage. The ordination shows stage-associated turnover in taxon occurrence, with fresh and pre-rotten fruit-days partly overlapping and post-rotten and degraded fruit-days shifting toward a more compositionally distinct region. The moderate stress value (stress = 0.221) indicates that the plot should be interpreted as a broad visual summary of compositional differences rather than an exact representation of all pairwise dissimilarities.

**Table S7.** Pairwise comparisons of Bray–Curtis multivariate dispersion among decomposition stages. **Model formula:** Distance to centroid ~ decomposition stage **Test:** Tukey HSD post-hoc comparison following beta disper analysis

| Comparison | Difference in distance to centroid | Lower 95% CI | Upper 95% CI | Adjusted p-value |
| --- | --- | --- | --- | --- |
| Pre-rotten - Fresh | 0.07763 | 0.00875 | 0.14650 | 0.02051 |
| Post-rotten - Fresh | 0.06069 | -0.00843 | 0.12981 | 0.10669 |
| Degraded - Fresh | -0.20375 | -0.33919 | -0.06831 | 0.00084 |
| Post-rotten - Pre-rotten | -0.01694 | -0.07347 | 0.03960 | 0.86314 |
| Degraded - Pre-rotten | -0.28138 | -0.41085 | -0.15191 | 0.0000006 |
| Degraded - Post-rotten | -0.26444 | -0.39404 | -0.13484 | 0.0000030 |

**Table S8.** Additive PERMANOVA models testing decomposition-stage and microhabitat effects on community composition. Additive PERMANOVA models testing the effects of decomposition stage and microhabitat on fruit-associated invertebrate community composition. Bray–Curtis dissimilarity was calculated from square-root-transformed peak relative abundance, whereas Jaccard dissimilarity was calculated from presence/absence data. Models were fitted using adonis2 with terms added sequentially and 999 permutations constrained within fruit identity to account for repeated sampling of the same fruits. Decomposition stage explained substantially more variation than microhabitat in both Bray–Curtis and Jaccard analyses, indicating that temporal succession during fruit decomposition was the dominant driver of community turnover. **Model formulas:** Bray–Curtis model: bray_peak ~ decomposition_stage + microhabitat Jaccard model: jaccard_pa ~ decomposition_stage + microhabitat Permutation structure: 999 permutations constrained within fruit identity

| Dissimilarity | Term | Df | Sum of squares | R<sup>2</sup> | F | p-value |
| --- | --- | --- | --- | --- | --- | --- |
| Bray–Curtis | Decomposition stage | 3 | 8.123 | 0.18491 | 9.2812 | 0.001 |
| Bray–Curtis | Microhabitat | 1 | 1.091 | 0.02483 | 3.7393 | 0.001 |
| Bray–Curtis | Residual | 119 | 34.718 | 0.79026 |  |  |
| Bray–Curtis | Total | 123 | 43.933 | 1.00000 |  |  |
| Jaccard | Decomposition stage | 3 | 6.543 | 0.14072 | 6.6436 | 0.001 |
| Jaccard | Microhabitat | 1 | 0.889 | 0.01912 | 2.7078 | 0.001 |
| Jaccard | Residual | 119 | 39.064 | 0.84017 |  |  |
| Jaccard | Total | 123 | 46.496 | 1.00000 |  |  |

**Table S9.**
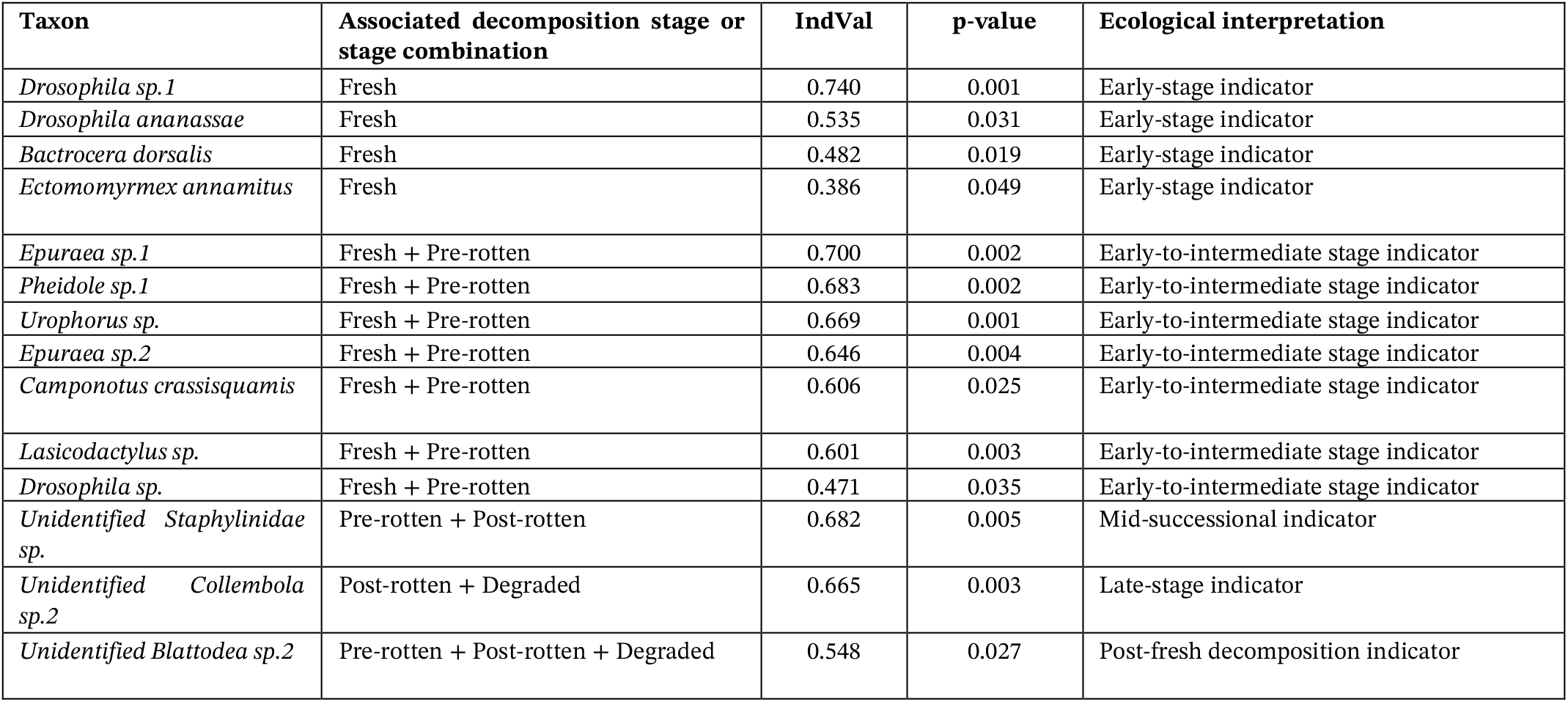
Significant indicator taxa associated with decomposition stages based on presence/absence data. Analysis method: Multilevel pattern analysis using IndVal.g Model grouping variable: Decomposition stage

**Table S10.**
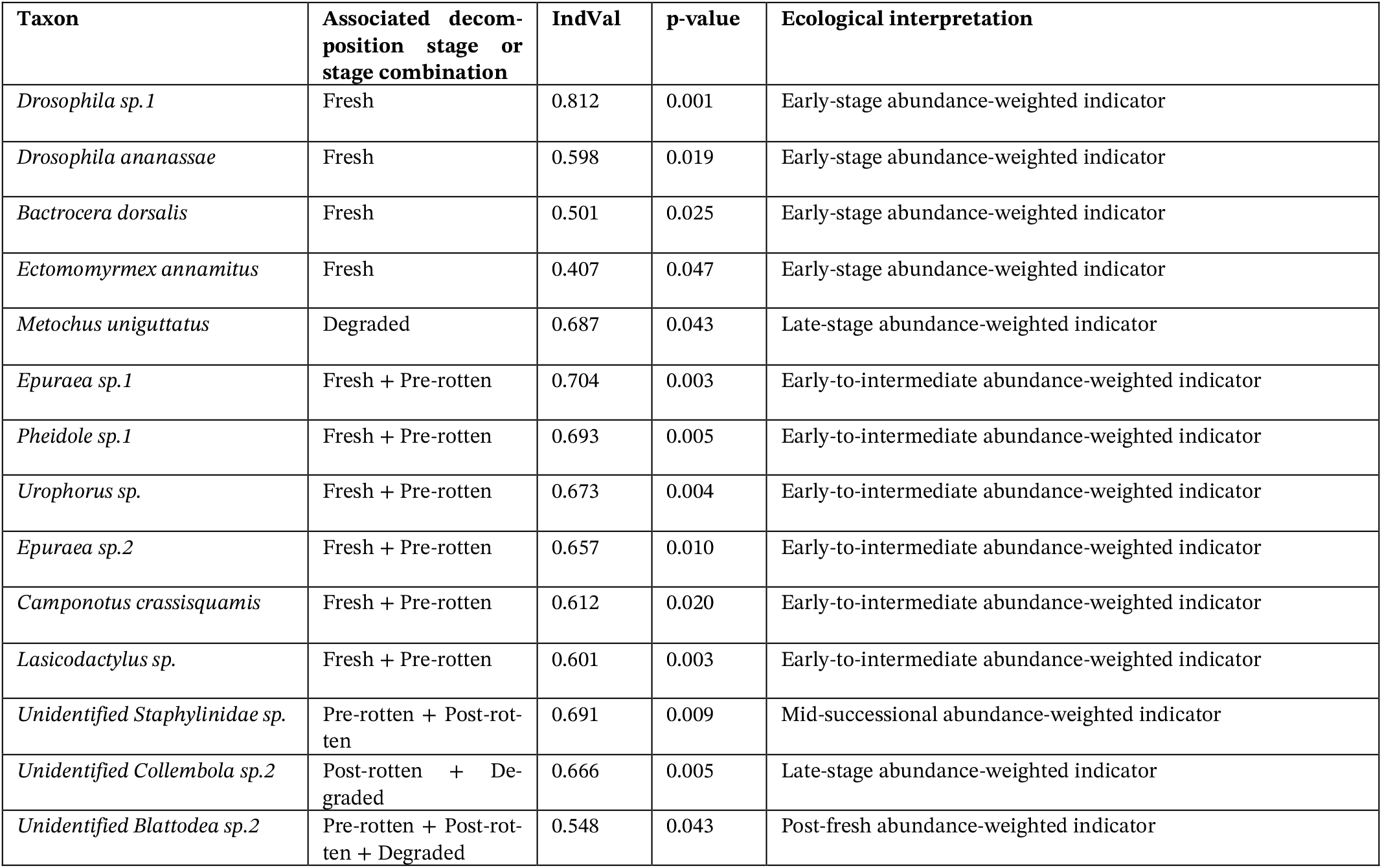
Significant indicator taxa associated with decomposition stages based on abundance-weighted data. Significant indicator taxa associated with decomposition stages of fallen guava fruits based on abundance-weighted data. Indicator taxa were identified using multilevel pattern analysis with the IndVal.g association function and 999 permutations. The analysis was performed on the square-root-transformed peak relative abundance community matrix. Fourteen of 44 taxa were significant at p < 0.05. Five taxa were associated with a single decomposition stage, eight taxa were associated with two-stage combinations, and one taxon was associated with a three-stage combination. Higher IndVal values indicate stronger abundance-weighted association with the corresponding decomposition stage or stage combination. Analysis method: Multilevel pattern analysis using IndVal.g

**Table S11.** Fixed-effect estimates from the Poisson GLMM testing the association between Pheidole sp.1 abundance and non-ant taxon richness. **Model formula:** Non-ant taxon richness ~ standardized Pheidole sp.1 peak abundance + microhabitat + decomposition stage + standardized temperature + standardized humidity + (1 | fruit identity)

| Predictor | Estimate | Std. Error | z value | p-value |
| --- | --- | --- | --- | --- |
| Intercept | 1.94561 | 0.24084 | 8.078 | 6.56e-16 |
| *Pheidole sp.1* peak abundance, standardized | -0.05726 | 0.03407 | -1.681 | 0.092852 |
| Microhabitat: Shaded | -0.06057 | 0.31035 | -0.195 | 0.845255 |
| Decomposition stage: Pre-rotten | 0.09562 | 0.11191 | 0.854 | 0.392896 |
| Decomposition stage: Post-rotten | -0.49542 | 0.13486 | -3.674 | 0.000239 |
| Decomposition stage: Degraded | -1.65482 | 0.23615 | -7.007 | 2.43e-12 |
| Temperature, standardized | 0.02544 | 0.08674 | 0.293 | 0.769303 |
| Humidity, standardized | -0.03306 | 0.05707 | -0.579 | 0.562402 |

**Table S12.** Fixed-effect estimates from the COM-Poisson GLMM testing the association between Pheidole sp.1 abundance and non-ant peak relative abundance. Fixed-effect estimates from the COM-Poisson generalized linear mixed model testing the relationship between Pheidole sp.1 peak abundance and non-ant peak relative abundance per fruit-day. The model used a COM-Poisson error distribution with a log link. Exposed microhabitat and fresh decomposition stage were used as reference categories. Fruit identity was included as a random intercept to account for repeated sampling of the same fruits through decomposition. The coefficient for standardized Pheidole sp.1 peak abundance was negative and marginal at p = 0.0568, suggesting a weak negative association with non-ant peak abundance, but not statistically significant at p < 0.05. **Model formula:** Non-ant peak relative abundance ~ standardized Pheidole sp.1 peak abundance + microhabitat + decomposition stage + standardized temperature + standardized humidity + (1 | fruit identity)

| Predictor | Estimate | Std. Error | z value | p-value |
| --- | --- | --- | --- | --- |
| Intercept | 4.14003 | 0.27652 | 14.972 | < 2e-16 |
| Pheidole sp.1 peak abundance, standardized | -0.13413 | 0.07041 | -1.905 | 0.0568 |
| Microhabitat: Shaded | 0.03137 | 0.32388 | 0.097 | 0.9228 |
| Decomposition stage: Pre-rotten | -0.11324 | 0.19104 | -0.593 | 0.5534 |
| Decomposition stage: Post-rotten | -1.07261 | 0.22479 | -4.772 | 1.83e-06 |
| Decomposition stage: Degraded | -2.78173 | 0.38078 | -7.305 | 2.77e-13 |
| Temperature, standardized | 0.06571 | 0.14076 | 0.467 | 0.6406 |
| Humidity, standardized | 0.19055 | 0.09861 | 1.932 | 0.0533 |

**Figure S5.**
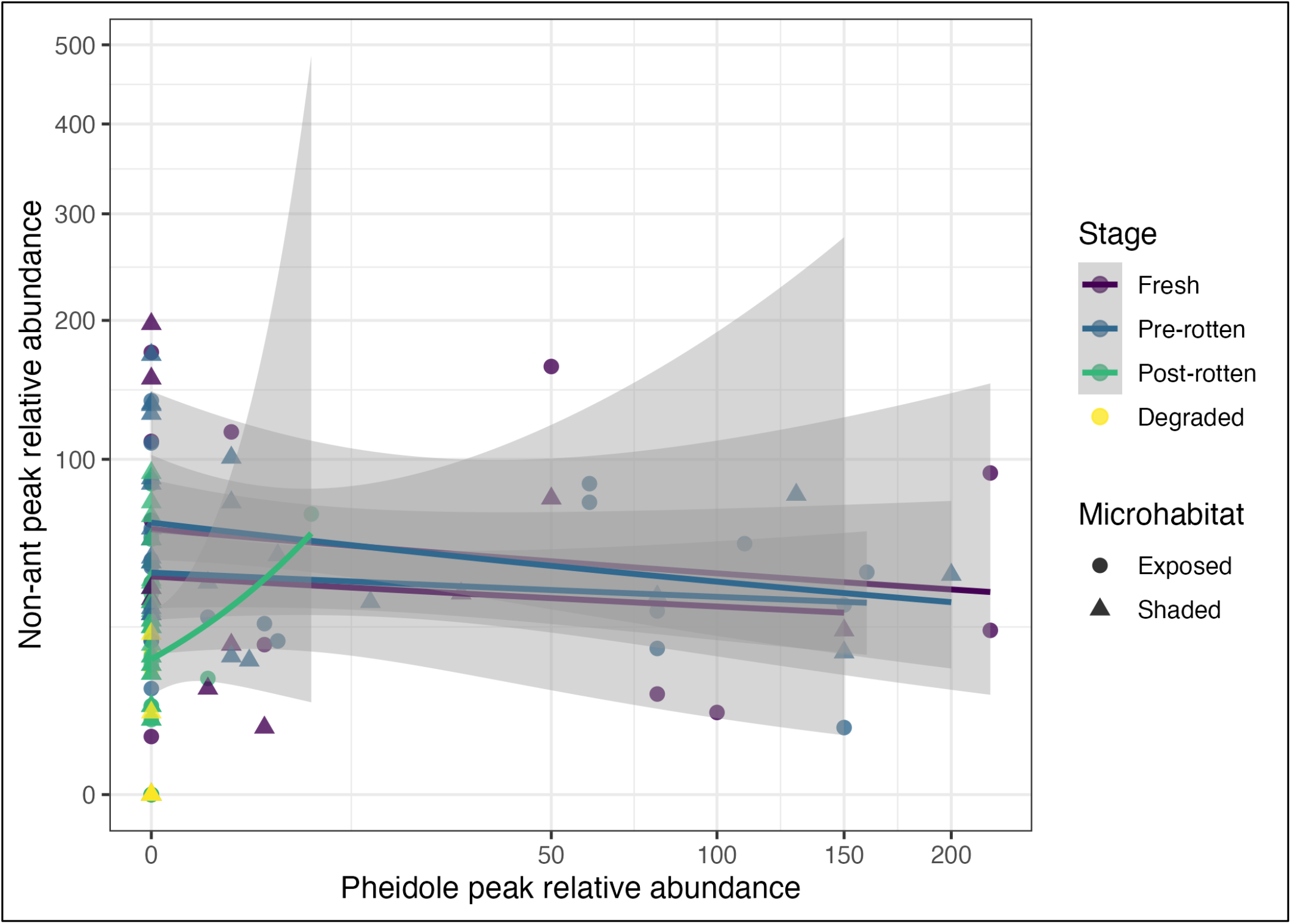
Relationship between Pheidole sp.1 activity and non-ant peak relative abundance. Scatterplot showing the relationship between Pheidole sp.1 peak relative abundance and non-ant peak relative abundance on fallen ripe Psidium guajava fruit-days. Points represent individual fruit-days, colours indicate decomposition stage, and shapes indicate microhabitat type: exposed or shaded. Lines show stage-specific fitted trends with confidence intervals. The plot provides an exploratory visualization of ant-associated patterns; although non-ant peak relative abundance tended to be lower at higher Pheidole sp.1 activity in some stages, mixed-model analyses showed only a marginal negative association after accounting for decomposition stage and other covariates. Therefore, the pattern should be interpreted as observational evidence of possible ant-associated variation in resource use, not as evidence of causal suppression.

**Figure S6.**
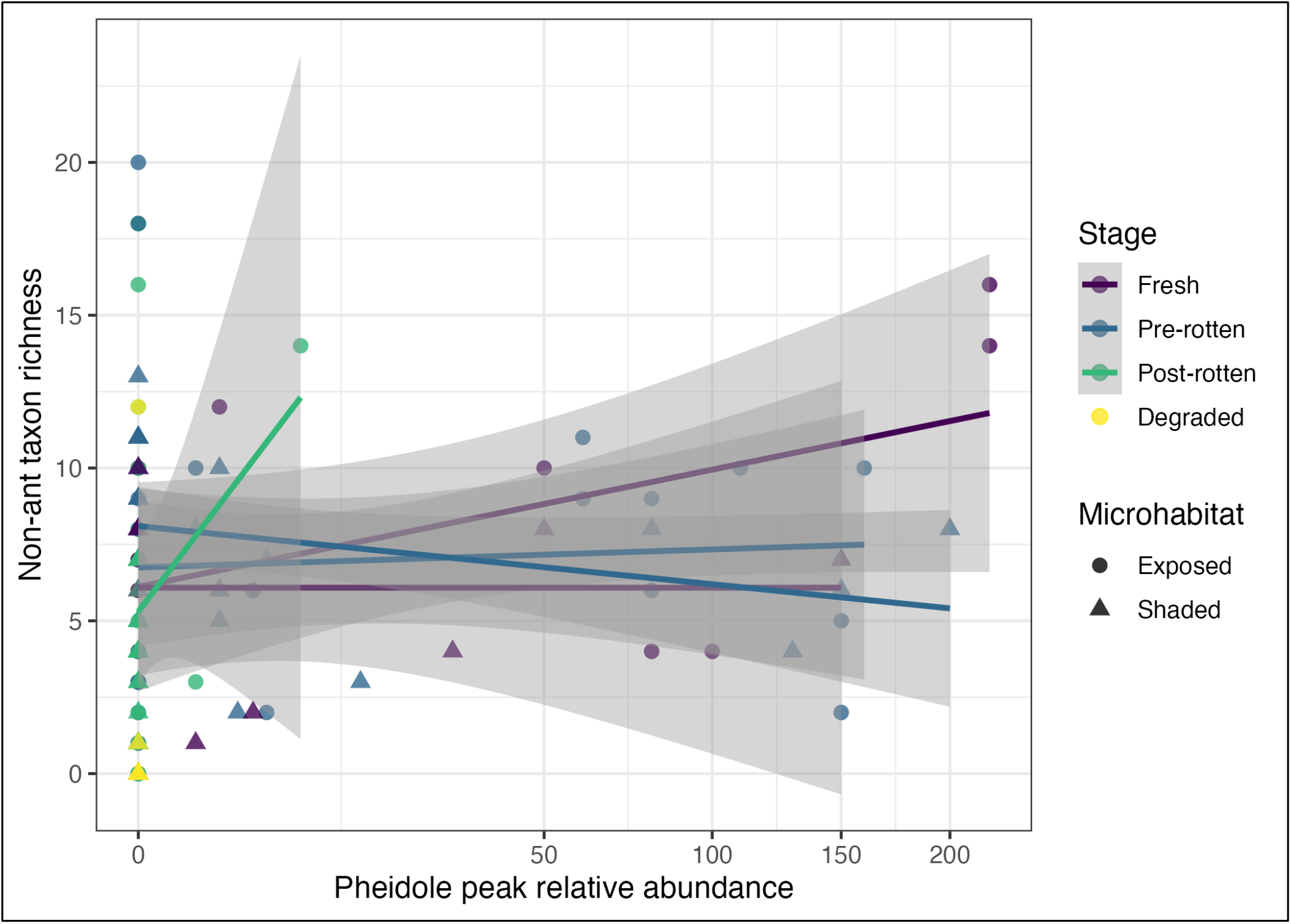
Relationship between Pheidole sp.1 activity and non-ant taxon richness. Scatterplot showing the relationship between Pheidole sp.1 peak relative abundance and non-ant taxon richness on fallen ripe Psidium guajava fruit-days. Points represent individual fruit-days, colours indicate decomposition stage, and shapes indicate microhabitat type: exposed or shaded. Lines show stage-specific fitted trends with confidence intervals. The figure provides an exploratory visualization of ant-associated patterns; mixed-model analysis showed a weak negative but non-significant association between Pheidole sp.1 peak relative abundance and non-ant richness after accounting for decomposition stage and other covariates. Therefore, the pattern should be interpreted as observational evidence of possible ant-associated variation in community structure, not as evidence of causal suppression.

**Table S13.** Fixed-effect estimates from the negative-binomial GLMM testing the association between Pheidole sp.1 abundance and Diptera peak relative abundance. **Model formula:** Diptera peak relative abundance ~ standardized Pheidole sp.1 peak abundance + microhabitat + decomposition stage + standardized temperature + standardized humidity + (1 | fruit identity)

| Predictor | Estimate | Std. Error | z value | p-value |
| --- | --- | --- | --- | --- |
| Intercept | 3.0410 | 0.4125 | 7.372 | 1.68e-13 |
| *Pheidole* sp.1 peak abundance, standardized | -0.2243 | 0.1489 | -1.506 | 0.13203 |
| Microhabitat: Shaded | 0.4004 | 0.4193 | 0.955 | 0.33962 |
| Decomposition stage: Pre-rotten | -1.1043 | 0.3704 | -2.981 | 0.00287 |
| Decomposition stage: Post-rotten | -2.7045 | 0.4637 | -5.832 | 5.47e-09 |
| Decomposition stage: Degraded | -5.3520 | 0.8995 | -5.950 | 2.68e-09 |
| Temperature, standardized | -0.2875 | 0.2546 | -1.129 | 0.25878 |
| Humidity, standardized | 0.4146 | 0.2244 | 1.847 | 0.06471 |

**Table S14.**
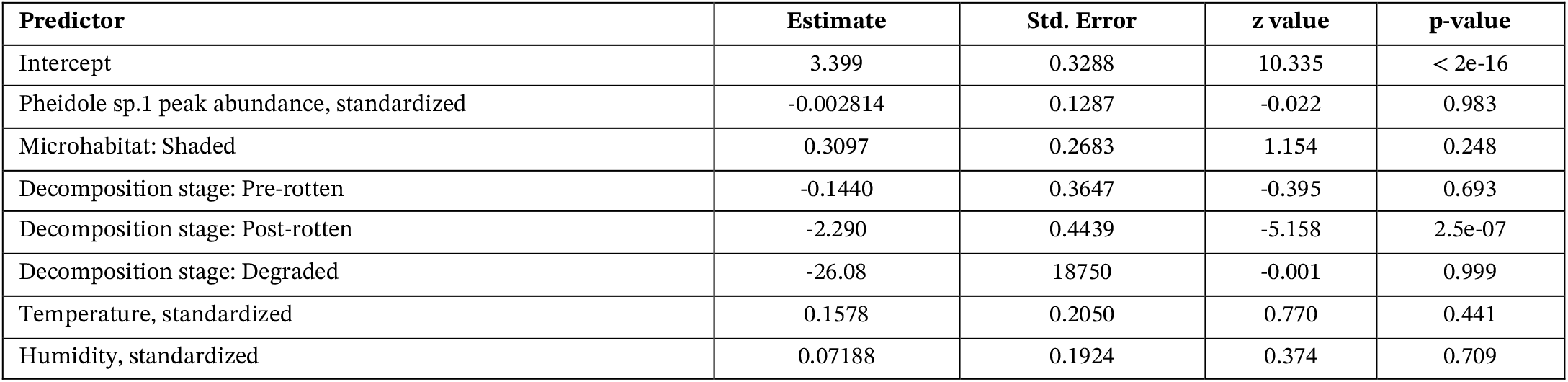
Fixed-effect estimates from the negative-binomial generalized linear mixed model testing the relationship between Pheidole sp.1 peak abundance and Coleoptera peak relative abundance per fruit-day. The model used a negative-binomial error distribution with a log link. Exposed microhabitat and fresh decomposition stage were used as reference categories. Fruit identity was included as a random intercept to account for repeated sampling of the same fruits through decomposition. Pheidole sp.1 peak abundance was not associated with Coleoptera peak abundance. Coleoptera peak abundance was significantly lower in the post-rotten stage than in the fresh stage. The degraded-stage estimate should be interpreted cautiously because the very large standard error suggests sparse or near-zero Coleoptera observations in degraded fruit-days. Model formula: Coleoptera peak relative abundance ~ standardized Pheidole sp.1 peak abundance + microhabitat + decomposition stage + standardized temperature + standardized humidity + (1 | fruit identity)

**Table S15.** Fixed-effect estimates from the negative-binomial GLMM testing the association between Pheidole sp.1 abundance and maggot/larval peak relative abundance. Fixed-effect estimates from the negative-binomial generalized linear mixed model testing the relationship between Pheidole sp.1 peak abundance and maggot/larval peak relative abundance per fruit-day. The model used a negative-binomial error distribution with a log link. Exposed microhabitat and fresh decomposition stage were used as reference categories. Fruit identity was included as a random intercept to account for repeated sampling of the same fruits through decomposition. Pheidole sp.1 peak abundance was not significantly associated with maggot/larval peak abundance. The degraded-stage estimate should be interpreted cautiously because the very large standard error indicates sparse or near-zero maggot/larval observations in degraded fruit-days. **Model formula:** Maggot/larval peak relative abundance ~ standardized Pheidole sp.1 peak abundance + microhabitat + decomposition stage + standardized temperature + standardized humidity + (1 | fruit identity)

| Predictor | Estimate | Std. Error | z value | p-value |
| --- | --- | --- | --- | --- |
| Intercept | 0.84369 | 1.21044 | 0.697 | 0.486 |
| Pheidole sp.1 peak abundance, standardized | -0.48930 | 0.34288 | -1.427 | 0.154 |
| Microhabitat: Shaded | -1.17692 | 1.21436 | -0.969 | 0.332 |
| Decomposition stage: Pre-rotten | -0.53468 | 1.01478 | -0.527 | 0.598 |
| Decomposition stage: Post-rotten | -1.30779 | 1.11343 | -1.175 | 0.240 |
| Decomposition stage: Degraded | -21.93657 | 7434.15406 | -0.003 | 0.998 |
| Temperature, standardized | 1.28289 | 0.83863 | 1.530 | 0.126 |
| Humidity, standardized | 0.01803 | 0.44335 | 0.041 | 0.968 |

**Figure S7.**
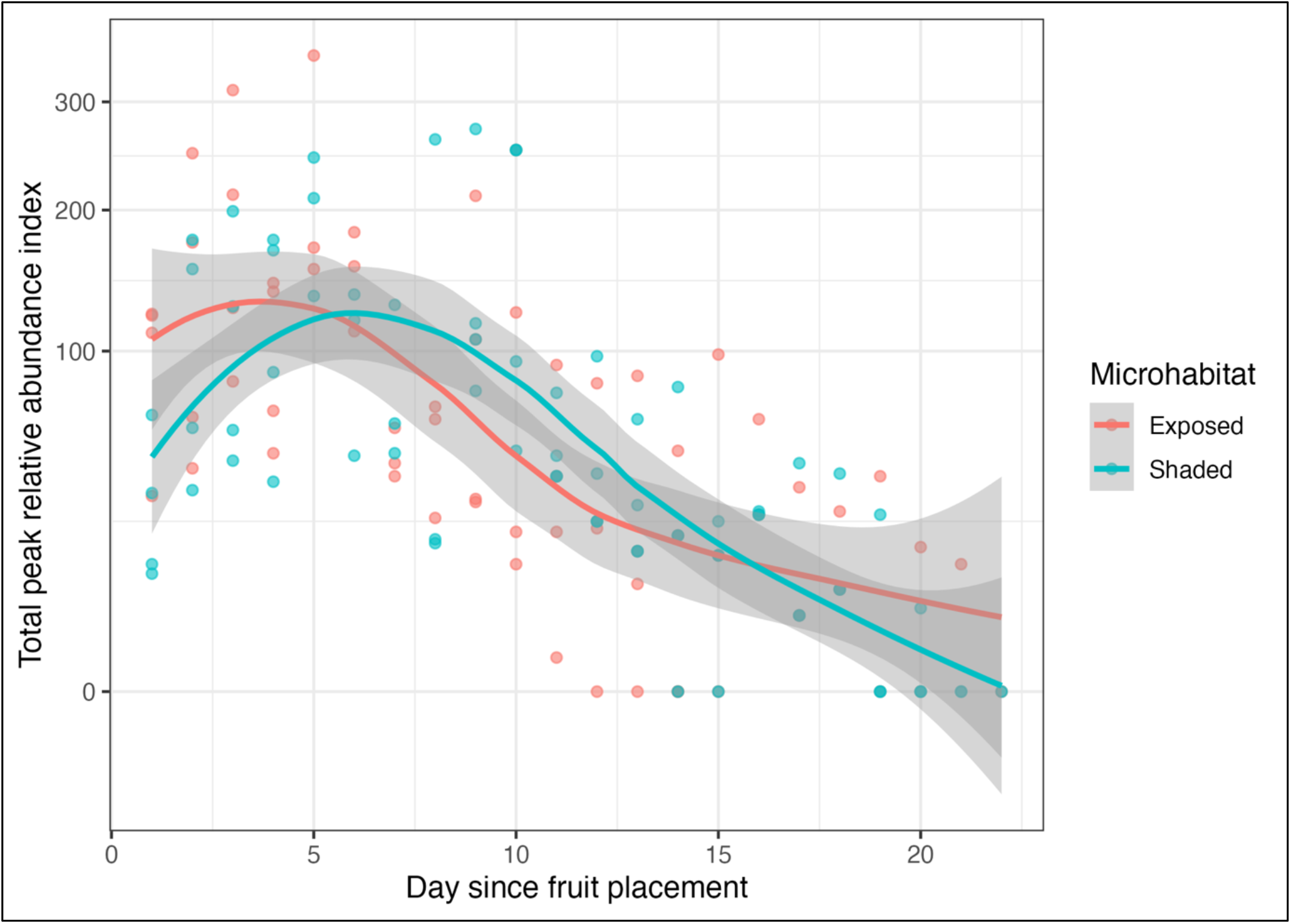
Day-wise change in total peak relative abundance on fallen ripe guava fruits. Total peak relative abundance index recorded on fallen ripe Psidium guajava fruits across days since fruit placement in exposed and shaded microhabitats. Points represent individual fruit-days, and lines show LOESS-smoothed trends with confidence intervals for each microhabitat. Total peak relative abundance was highest during early decomposition, particularly during the fresh and pre-rotten stages, and declined strongly as fruits progressed toward the post-rotten and degraded stages. The plot is used as a descriptive visualization of continuous day-wise succession and supports the stage-based analyses showing reduced invertebrate activity during later decomposition.

**Figure S8.**
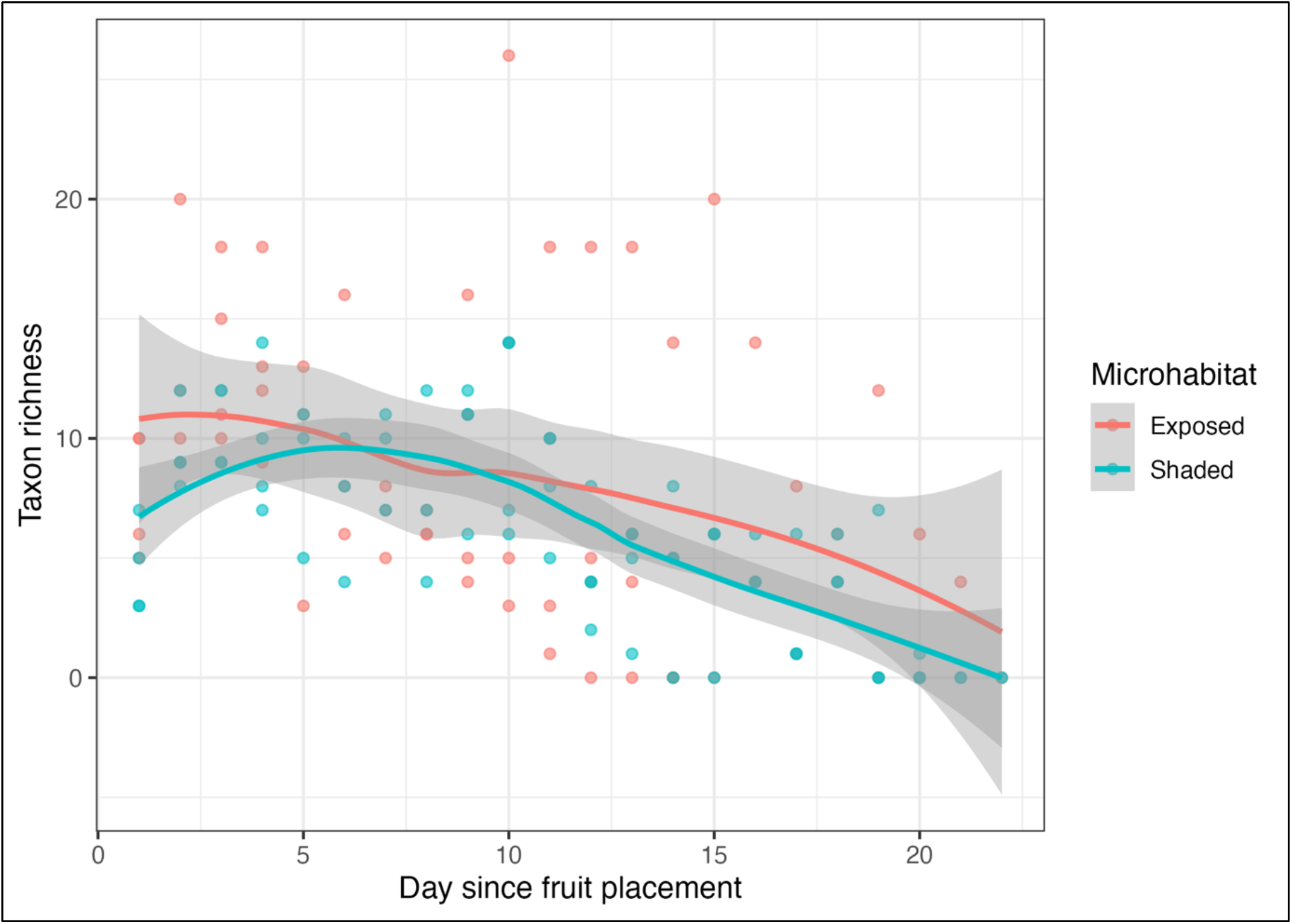
Day-wise change in taxon richness on fallen ripe guava fruits. Taxon richness recorded on fallen ripe Psidium guajava fruits across days since fruit placement in exposed and shaded microhabitats. Points represent individual fruit-days, and lines show LOESS-smoothed trends with confidence intervals for each microhabitat. Richness remained relatively high during the early days of decomposition and declined progressively as fruits became post-rotten and degraded. The plot is used as a descriptive visualization of continuous day-wise succession and supports the stage-based analyses showing a strong decline in richness during later decomposition.

